# Biohybrid Microswimmers Against Bacterial Infections

**DOI:** 10.1101/2021.05.10.443410

**Authors:** I. S. Shchelik, J. V. D. Molino, K. Gademann

## Abstract

Biohybrid microswimmers exploit the natural abilities of motile microorganisms *e.g.* in releasing cargo on-demand with high spatial and temporal control. However, using such engineered swarms to deliver antibiotics addressing bacterial infections has not yet been realized. In the present study, a design strategy for biohybrid microswimmers is reported, which features the covalent attachment of antibiotics to the motile green algae C*hlamydomonas reinhardtii* via a photo-cleavable linker. The surface engineering of the algae does not rely on genetic manipulations, proceeds with high efficiency, does not impair the viability or phototactic ability of microalgae, and allows for caging of the antibiotic on the surface for subsequent release via external stimuli. Two different antibiotic classes have been separately utilized, which result in activity against both gram-positive and gram-negative strains. Guiding the biohybrid microswimmers by an external beacon, and on-demand delivery of the drugs by light with high spatial and temporal control, allowed for strong inhibition of bacterial growth *in vivo*. This efficient strategy could potentially allow for the selective treatment of bacterial infections by engineered algal microrobots with high precision in space and time. Overall, this work presents an operationally simple production of biohybrid microswimmers loaded with antibiotic cargo to combat bacterial infections precisely delivered in three-dimensional space.

---

Microrobots have recently received strong attention in various fields, due to their excellent ability to conduct microscale tasks with surprisingly high efficiency via dedicated movements in response to specific stimuli. However, the development of reliable and efficient actuators, power sources, or sensors on a micrometer scale remain as challenges.^[1]^ Most synthetic microdevices involve external propulsion mechanisms, such as chemically powered motors^[2,3]^ driven by hydrogen peroxide,^[4–6]^ glucose^[7,8]^ or acidic solution,^[9]^ involving metal or enzyme catalysts.^[10]^ Different examples of propulsion mechanisms are based on fuel-free sources such as magnetic fields,^[11]^ visible light^[12–15]^, ultrasound^[16,17]^ or electric fields.^[18,19]^ Nevertheless, those micromotors imply significant challenges in compatibility with high ionic strength media,^[20]^ movement in biological fluids,^[21–23]^ production of toxic by-products, and having limited speed, power or directionality.^[24,25]^ On the other hand, biological cells have evolved natural movement capabilities and demonstrate efficient propulsion on the microscale. These organisms can respond to environmental stimuli such as chemicals (chemotaxis),^[26]^ light (phototaxis)^[27,28]^ or pH gradients (pH-taxis)^[29]^ through integrated sensing and conversion of chemical energy into mechanical action. These cellular functionalities have already been utilized for the development of novel bio-hydrid microswimmers,^[30,31]^ in which bacteria are commonly used as a self-powered micro-actuators responding to different stimuli.^[32,33,34]^ Bacteria have been used for carrying microbeads,^[35–42]^ microtubes,^[43]^ or double emulsions,^[44]^ driving rotatory motors,^[35,45]^ moving solid structure,^[46,47]^ or transportation of live red blood cells.^[48]^ However, bacteria-based biohybrid systems face multiple issues including potential pathogenicity or immunogenicity, rapid growth rate, and their potential to develop antibiotic resistance.^[49]^

Biohybrid systems derived from other motile cell types can complement bacterial approaches and benefit from (1) improved biocompatibility, (2) slower growth, (3) better mobility in the physiologically relevant media, and (4) absence of known toxins. For example, sperm micromotors displayed high potential in moving against blood flow or in oviduct fluid and viscoelastic media while serving as a cargo transporter.^[50–53]^ In this regard, microalgae were investigated as a microrobotic platform for on-demand cargo delivery^[27,54,55]^, imaging-guided therapy^[56]^ or on-cell catalysis.^[57]^ *Chlamydomonas reinhardtii* is a biflagellate unicellular green microalgae with 10 μm in diameter possessing phototaxis properties and autofluorescence, which allows noninvasive tracking of the cells without additional fluorescent biomarkers.^[58]^ Additionally, the negatively charged *C. reinhardtii* cell wall was used for the modification of interacting surfaces by electrostatic interactions. For example, the coating of the microalgae surface was achieved with chitosan,^[27]^ 4-hydroxyproline (4-HP)-rich polypeptides,^[59,60]^ or magnetic microbeads.^[61]^ Covalent attachment of artificial metalloenzymes^[57]^ and antibiotics^[62]^ were recently reported by our group. This strategy allows to reach high loading efficacy and does not suffer from the loss of the payload from the cell surface. *C. reinhardtii* is known as a phototactic green microalgae, which swim steadily towards or away from the light stimulus due to optical receptors.^[63]^ The highest phototactic response is observed for the microalgae at exponential growth phase and it happens in a matter of seconds reaching the linear stage within 1-3 minutes.^[64]^ Conclusively, *C. reinhardtii* phototaxis is a highly desirable trait since it makes the motion of the produced biohybrids controllable and steerable.

In this work, we leverage the previously developed surface engineering of microalgae to obtain biohybrids with phototactic activity, providing three-dimensional localization of antibiotic activity in the region of interest. Two antibiotics with different mode of action - vancomycin (VAN) and ciprofloxacin (CIP) were attached to the surface of *C. reinhardtii* by covalent binding without any adverse effects on motility and phototactic properties of the biohybrid microalgae. Moreover, we performed on-demand delivery of the drug and its release by UV-irradiation at λ = 365 nm leading to successful bacterial growth inhibition. Overall, this work presents an operationally simple production of biohybrid microswimmers loaded with antibiotic cargo to combat bacterial infections precisely delivered in three-dimensional space.

Strategies for covalent bond formation between the cargo and the surface of living cells have utilized the presence of hydroxyproline-rich heteropolysaccharides in *Chlamydomonas*.^[65]^ A two-step approach was performed for coating, including the functionalization of the cell surface with an NHS-dibenzocyclooctyne anchor (DBCO) as a first step followed by the antibiotic conjugation with an azide derivative of the drug on the second step *via* biorthogonal click chemistry (Figure 1, A). The scope of antibiotics used for the attachment to microalgae surface includes two widely used antibiotics with different mode of action (Figure 1, B). Moreover, the successful functionalization of the microalgae surface was demonstrated by confocal laser scanning microscopy where coating of the algal cell was observed after cell wall modification with a fluorescein-containing vancomycin derivative (Figure 1, C).^[62]^

**Figure 1.**
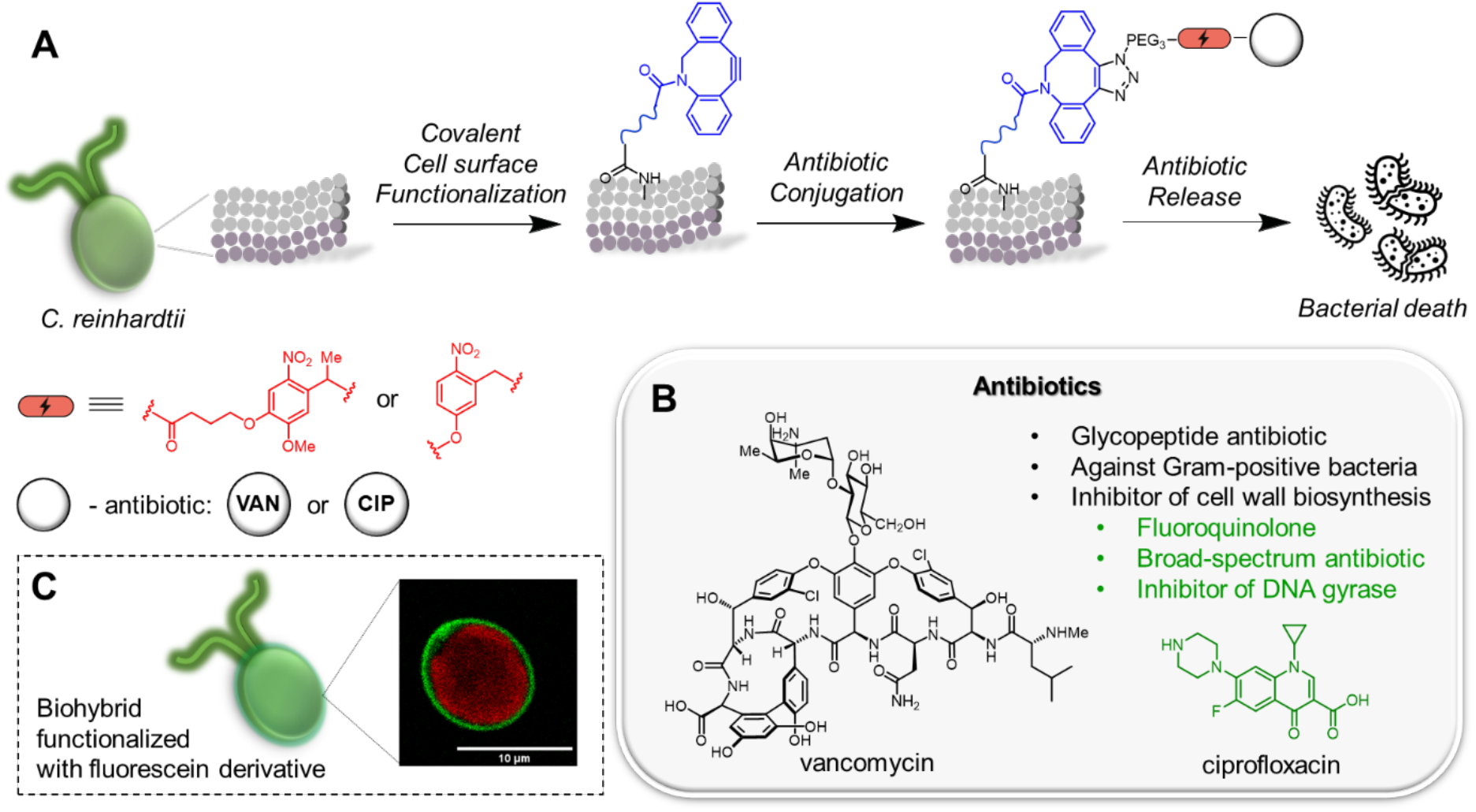
Biohybrid *C. reinhardtii* microswimmers production. A) Schematics of microalgae surface functionalization leading to the production of biohybrid *C. reinhardtii* for the treatment of bacterial infections. B) Target antibiotics for the functionalization of the microalgae surface. C) Confocal laser scanning microscopy image of biohybrid microalgae functionalized with DBCO (0.30 mM) and fluorescein-containing vancomycin derivative (0.2 mM).^[62]^

To produce the desired antibacterial biohybrids, several steps need to be carried out. First, the target antibiotics must contain: a) the moiety for the covalent binding to the surface of microalgae, and b) the photocleavable linker to control release of an antibiotic via external stimuli. Secondly, the resulting biohybrids microswimmers need to (1) remain motile, (2) maintain phototaxis properties, and (3) retain the target drug delivery competence, such as antibacterial activity after the antibiotic release.

For the conjugation of the antibiotics to the microalgae surface, an azide group with a photocleavable linker was introduced into the target drugs. The synthesis of the vancomycin derivative via three synthetic steps was previously reported by our group (Figure 2, A).^[62]^ Ciprofloxacin was modified using similar synthetic approach: The ethyl ester of ciprofloxacin (**3)** was first reacted with the MOM-protected nitrobenzyl linker **4** via a reductive amination reaction following by MOM group deprotection resulting in compound **5**. Next, the incorporation of azide-containing linker was carried out via an S*N*2 reaction followed by the hydrolysis of ester leading to the desired product **6** over two steps (Figure 2, B).

**Figure 2.**
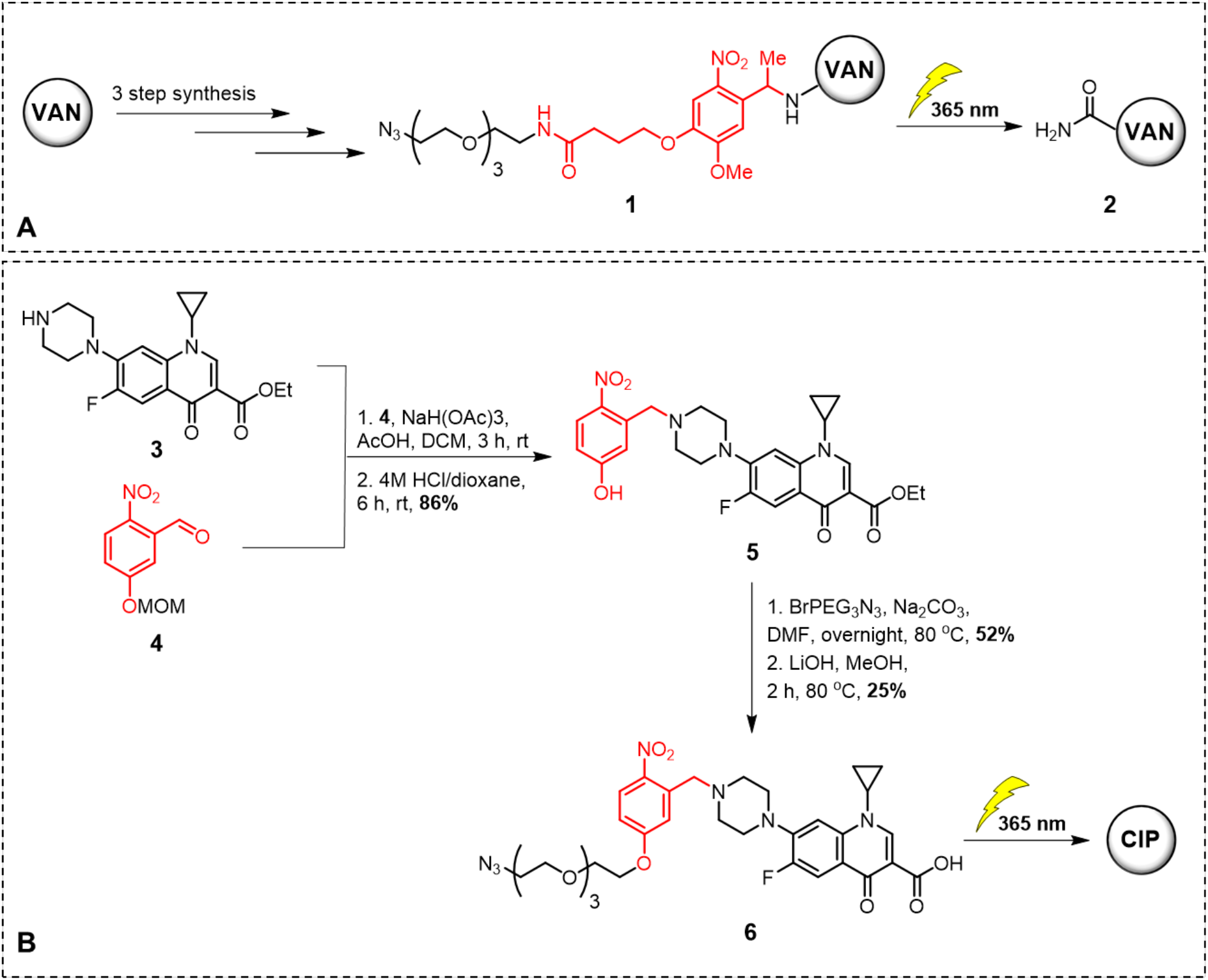
Syntheses of derivatives used for the functionalization of *C. reinhardtii* surface to produce biohybrid and the active molecules released after the UV-irradiation at λ = 365 nm. A) Synthesis of vancomycin derivative **1** and release of amide derivative of vancomycin (VAN-NH_2_, **2)**.^[62]^ B) Synthesis of ciprofloxacin derivative **6** and released active ciprofloxacin (CIP) after the UV-irradiation at λ = 365 nm.

The general photochemistry of the obtained antibiotic derivatives was studied next. Previously, it has been shown that the amide derivative VAN-NH_2_ **2** derives from the photocleavage of **1**.^[62]^ The photocleavage efficacy reaches 70% after 5 min of UV-irradiation at λ = 365 nm of vancomycin derivative **2** solution.^[62]^ Ciprofloxacin was characterized as product after the photocleavage of derivative **6** by UHPLC analysis (Figure S7, Supporting Information). The conversion from **6** to CIP by UV-irradiation was determined analyzing the formation of CIP by UHPLC-MS measurements in SIM mode (applied for the limit detection units) using a commercially available antibiotic as analytical standard. A maximum of 78% conversion was observed after 3 min of irradiation time, remaining constant until 5 min. This time length was chosen for the drug release experiments. Longer irradiation time caused drug decomposition (Figure S10, Supporting Information). Since our previous study^[62]^ demonstrated no impact on microalgae viability, bacterial growth and sufficient drug release after 5 min of UV-irradiation. Next, the coating of the microalgae with the active compounds was evaluated. Microalgae (1.5 × 10^6^ cells/mL) were incubated with three different concentrations of DBCO and antibiotics to evaluate the binding capability to the cells. All experiments for the cell surface functionalization were carried out in diluted microalgae growth medium (tris-acetate-phosphate (TAP) TAP:water in 1:10 ratio) to reduce stress for the cells due to the environmental change. The cells were first washed with dilute TAP medium and treated with DBCO anchor at different concentrations (0.05 mM, 0.1 mM and 0.2 mM). To ensure the availability of free cyclooctene-binding sites, a 10:1 ratio of DBCO:antibiotic was used. Then, DBCO-coated microalgae were washed and treated with either VAN-photoN_3_ **2** or CIP-photoN_3_ **6** at concentrations 0.005 mM, 0.01 mM or 0.02 mM respectively to DBCO concentrations leading to the desired biohybrids. The coating efficiency of biohybrid surface with antibiotics was calculated from the unreacted antibiotic washed out after the second incubation step. The coating yields of ≈90-95% were calculated after the functionalization of microalgae with antibiotics **2** or **6** (Figure S13, Supporting Information). The labelling efficacy with vancomycin derivative **2** have improved compared to the previously reported protocol (60-70%),^[62]^ which could be an effect of different solvents, concentrations or conditions during the incubation step. Next, for the evaluation of the light-triggered drug release, the biohybrids were UV-irradiated at λ = 365 nm for 5 min. The cells were washed with dilute TAP medium (6 times) and the supernatant was collected to analyzed released antibiotics by UHPLC (SIM mode). The release efficacy (%) was calculated from the amount of the drugs loaded on the cells and reached 5-10%. The lower yields can be explained by either the proximity of the drugs to the surface of microalgae, which could shade the linker from the light, or absorption of UV-light by microalgae,^[66]^ thereby influencing on cleavage efficacy. Nevertheless, the measured concentrations of released VAN-NH_2_ **3** (0.27 μg mL^−1^) and CIP (0.15 μg mL^−1^) were higher that the MIC value determined against *B. subtilis* and *E. coli*, respectively (Table S2, Supporting Information) when using the lowest loading concentrations of DBCO and the drug. Unfortunately, the highest tested loading concentration for both VAN-photoN_3_ **2** and CIP-photoN_3_ **6** (0.02 mM) is not sufficient to kill *S. aureus* (MRSA) after the photocleavage and release of the active drug (Figure 3). Thereby, the double increase in the concentrations of DBCO and antibiotic (0.4 mM and 0.04 mM, respectively) for the functionalization of microalgae surface was chosen to deliver a biohybrid microswimmer with activity against *S. aureus*.

**Figure 3.**
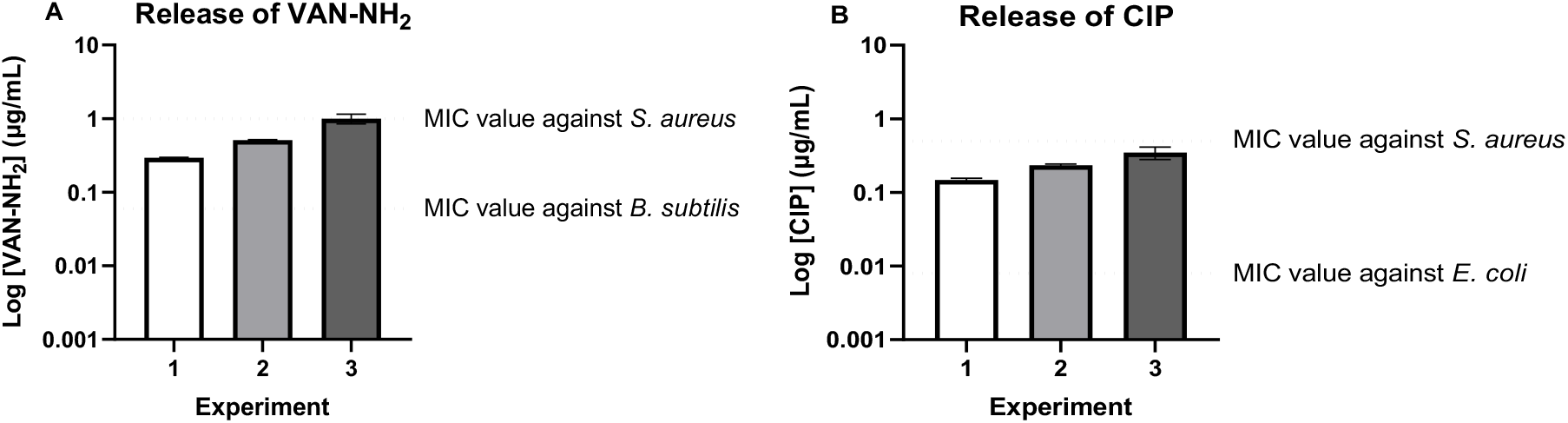
Release efficacy of antibiotics from the surface of *C. reinhardtii* after the UV-irradiation for 5 min at λ = 365 nm. The algae were modified with DBCO (Exp. 1: 0.05 mM; Exp. 2: 0.1 mM; Exp. 3: 0.2 mM) and either with VAN-photoN_3_ **2** (Exp. 1: 0.005 mM; Exp. 2: 0.01 mM; Exp. 3: 0.02 mM) or with CIP-photoN_3_ **6** (1: 0.005 mM; 2: 0.01 mM; 3: 0.02 mM). A) Release of VAN-NH_2_. B) Release of CIP. The dot lines correspond to MIC results against *S. aureus* ATCC 43300 (MRSA), *B. subtilis* ATCC 6633, *E. coli* ATCC 25922. Data are represented mean value ± SD (n = 3).

Next, we studied the motility of the antibiotic-loaded microswimmers by a series of experiments. The 2D swimming trajectory was recorded by a confocal laser scanning microscopy and analyzed using the software FIJI, and the plugin TrackMate (Figure 4, A). First, the swimming of biohybrid microalgae featuring fluorescein derivative of vancomycin coating was clearly observed (Movie S1, Supporting Information). Movie analyses revealed that the biohybrids remain motile in the presence of cargo on their surfaces. Moreover, the velocity of wild type and biohybrid microalgae did not show significant differences: Wild type microalgae displayed a median swimming velocity of 63.2 μm s^−1^, where the median swimming speed of biohybrid microalgae was 78.4 μm s^−1^ and 61.5 μm s^−1^ after the surface modification with VAN-photoN_3_ **3** and CIP-photoN_3_ **6**, respectively (Figure 4, B). The modification of microalgae did not impact on the speed of the cells and correspond well to the earlier published data.^[67,68]^

**Figure 4.**
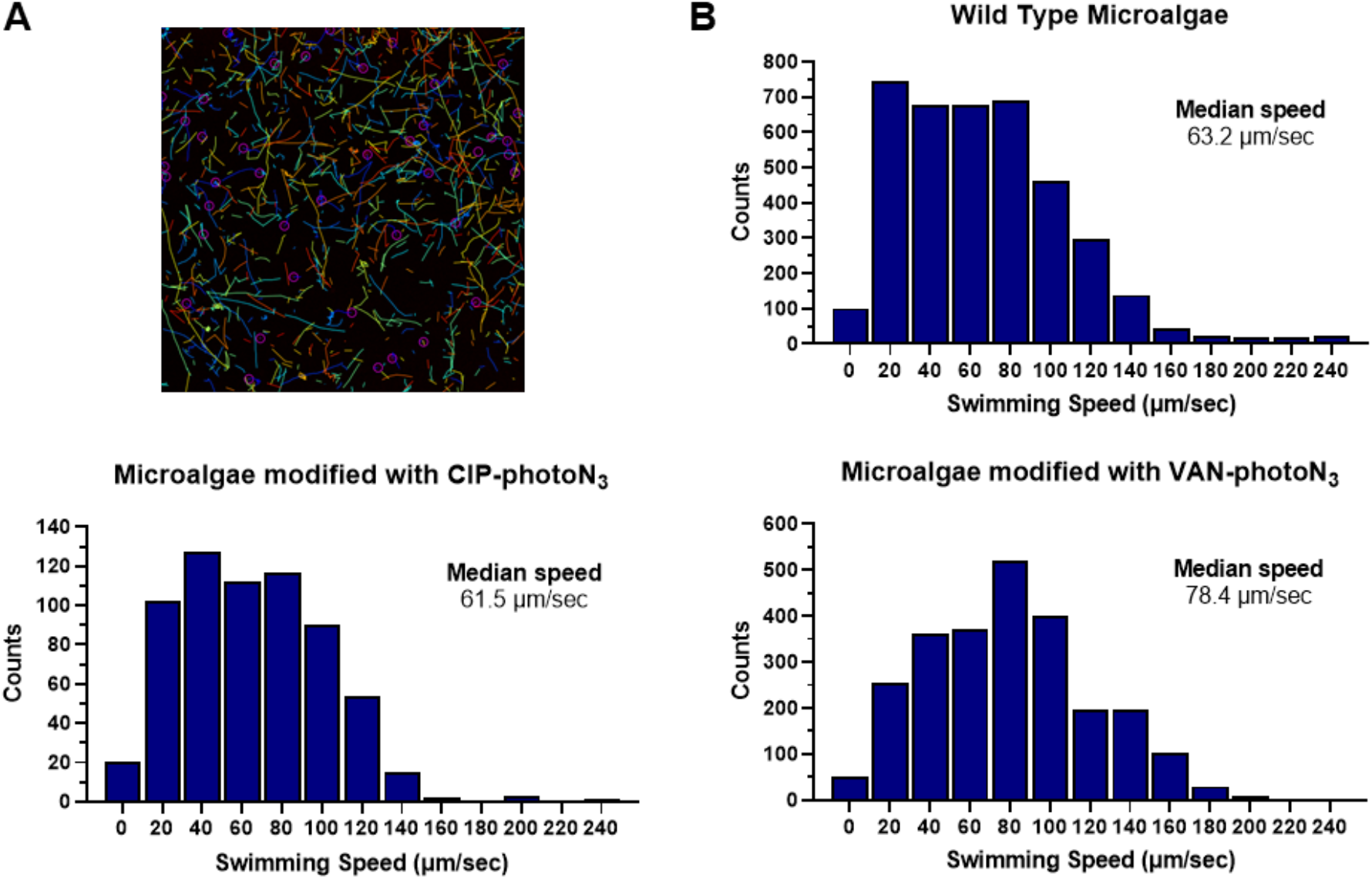
Velocity characterization of biohybrid *C. reinhardtii* microswimmers. A) An example of 2D swimming trajectories identified by TrackMate FIJI. B) 2D swimming speed analysis of bare microalgae and microalgae modified with either VAN-photoN_3_ or CIP-photoN_3_.

We investigated next whether our functionalization strategy by different antibiotics impairs the microalgae phototaxis properties. First, the microalgae were functionalized with DBCO (0.4 mM) and either VAN-photoN_3_ **2** or CIP-photoN_3_ **6** (0.04 mM). The concentration was chosen considering that the produced biohybrid microswimmers were active against *S. aureus*. Phototaxis behavior of wild type and biohybrid microswimmers were demonstrated in a cell culture well with a diameter of 2.5 cm. The wells filled with either wild type or biohybrid microalgae in water were illuminated from the right side for 5 min (Figure 5, A). The migration of wild type cells and their accumulation was observed on the opposite side of the light source. At this concentration, the modified microalgae did not show signs of phototaxis (Figure S17, Movie S2-S3, Supporting Information). Therefore, the microalgae were functionalized with a lower concentration of DBCO (0.2 mM) and antibiotics **2** or **6** (0.02 mM). Under these conditions, the migration and accumulation of both wild type and biohybrids were successfully observed on the left side of the well after light exposure (Figure 5, B; Movie S4-S5, Supporting Information). Quantification of microalgae cells at each side of the wells were performed to establish the difference in cellular density in each position after cells migration. The cells were collected from the left and right sides of the wells and the number was quantified by measuring OD750 using a plate reader (utilizing the calibration curve) and compared at 0 and 5 min time points (Figure 5, C). A clear difference in number of microalgae after migration was observed for wild type cells (ca. 80%) and for biohybrid cells (ca. 74/75 %). These data demonstrate the mildness of the functionalization treatment, in which the biohybrid microalgae retains the same capacity to be steered away from the light stimulus in a fast and controlled manner as the wild type cells. Moreover, introduction of antibiotics at a biologically useful concentration ≤ 0.02 mM did not reduce phototactic response of biohybrid microalgae. In addition, cells collected from different sides were UV-irradiated for 5 min at λ = 365 nm and the released antibiotics were analyzed by UHPLC (SIM mode) (Figure 5, D). Collectively, the obtained results demonstrate that it is possible to direct biohybrid microswimmers to the target location of bacterial accumulation and release sufficient amounts of antibiotic to inhibit bacterial growth for both VAN and CIP. Moreover, we could not detect any loss of antibiotics from the surface of the microswimmers after at least 2 h (Figure 5, D).

**Figure 5.**
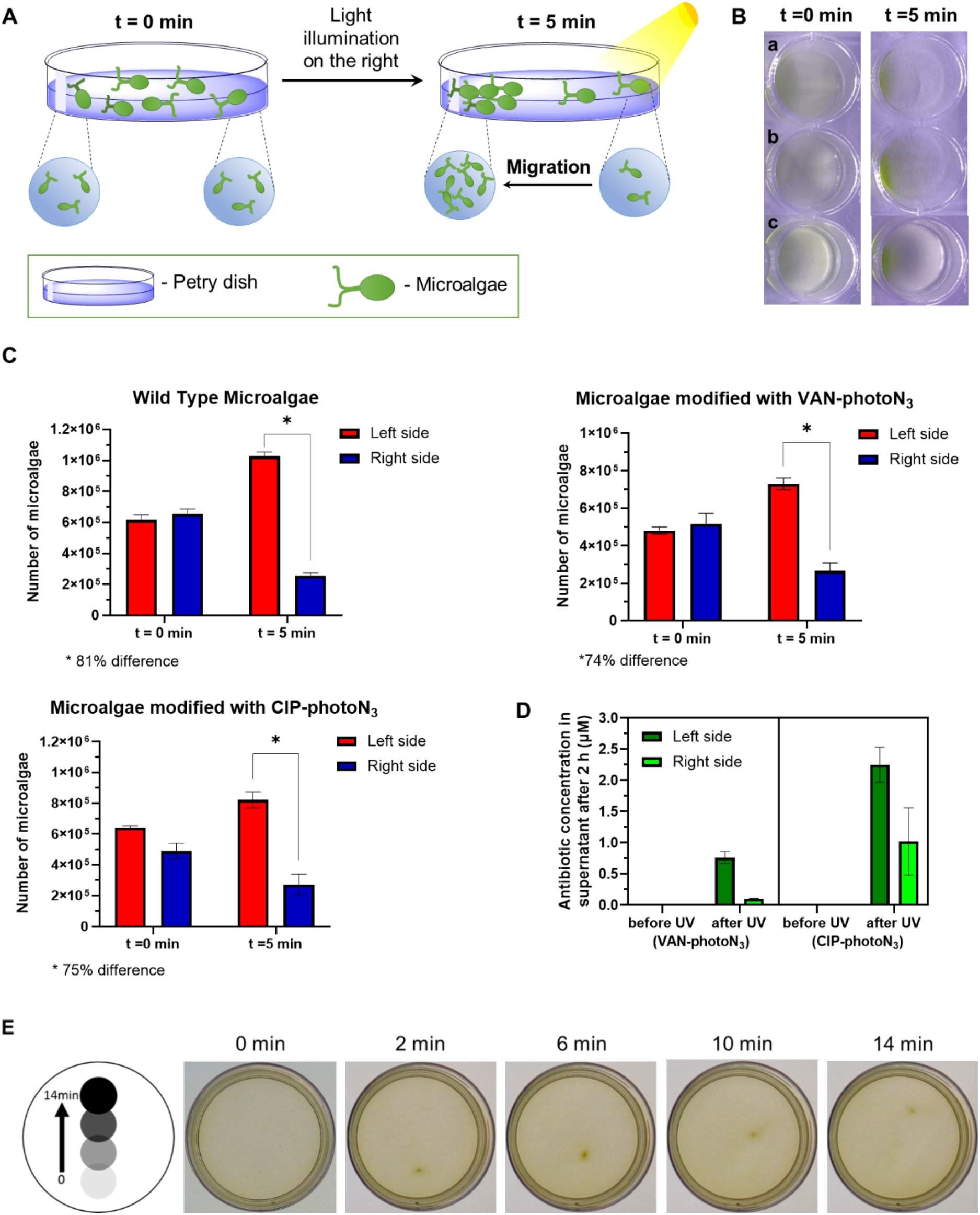
Phototaxis of biohybrid *C. reinhardtii* microswimmers. **A**) Schematic representation of light-driven steering of microalgae under visible light. The right side of the Petri dish was illuminated for 5 min to steer microalgae to the left. **B**) Images of microalgae before visible light exposure and after 5 minutes under visible light applied from the right side. (a) wild type microalgae (b) microalgae modified with VAN-photoN_3_ (c) microalgae modified with CIP-photoN_3_. **C**) Quantification of microalgae number at each end of the Petri dish. **D**) Drug release from microalgae surface collected from different sides of Petri dish before UV irradiation and after UV-irradiation for 5 min at λ = 365 nm. Microalgae modified with VAN-photoN_3_ – lift side; microalgae modified with CIP-photoN_3_ – right side. Data are represented mean value ± SD (n = 3). **E**) Microalgae swarm moving experiment. The cells swarm around the black dot area displayed below the well and follow the black dot movement. The video used for displaying the dot is Movie S6, Supporting Information.

To evaluate the effect of a high concentration of antibiotics on microalgae, a viability experiment was carried out. Microalgae (1.5 × 10^6^ cells/mL) were treated with antibiotics **2** or **6** at different concentrations (0.04 mM – 0.0025 mM) and the cell growth was recorded by measuring optical density OD_750_ using a plate reader for 4 days. No visible effect on microalgae growth was observed for the cells treated with the highest antibiotic concentration (Figure S20-21, Supporting Information). It was previously demonstrated that the exposure of the microalgae to high concentration of antibiotic induces the loss of flagella.^[60]^ As growth was not impaired, we hypothesize that loss of flagella function could be the explanation for the reduced phototaxis response of microalgae modified with 0.04 mM of antibiotics in our experiments.

To investigate the possibility to control the microalgae movement as a swarm, the following experiment was carried out. The microalgae (3 × 10^6^ cells/mL) were placed into a petri dish and the system was illuminated with a screen displaying a dark dot (Movie S6, Supporting Information). The designed dot was placed at the bottom side of the plate and then slowly moved to the top during 12 min. The cells accumulated on the dot area during the first two minutes and were steered to the top by moving the dot to the top of the dish (Figure 5, E). The calculated speed of microalgae swarm was 25 μm/sec, reflecting the dot speed. By this experiment, we demonstrated the direct control of microalgae movement using phototactic properties of the cells.

Next, we moved to the investigation of antibacterial activity of our biohybrid microswimmers against several bacterial strains. One of the most widespread and virulent nosocomial pathogens is methicillin-resistant *Staphylococcus aureus* (MRSA).^[69]^ It is a major cause of invasive disease among hospitalized patients and responsible for skin and soft tissue infections affecting up to one million people every year in the United States alone.^[70]^ Therefore, we decided that a *S. aureus* MRSA strain would be an excellent choice to test the antibacterial activity of our biohybrid modified with vancomycin derivative. Moreover, since we coated the microalgae with broad-spectrum antibiotics - ciprofloxacin, Gram-negative *E. coli* strains were also included into our study. The microalgae were modified with VAN-photoN_3_ (**2**, 0.02 mM) or CIP-photoN_3_ (**6**, 0.02 mM) according to the general protocol (Figure 1, A). Two samples with modified microalgae were combined when tested against *S. aureus* to obtain the doubled amount of the antibiotics in the solution after the release. The biohybrids were added into a suspension of *S. aureus* or *E. coli* in 24-well plate. The system was UV-irradiated at λ = 365 nm for 5 min. As a negative control experiment, the biohybrids were added after irradiation. The samples were incubated for 18 h, and the bacterial growth was recorded by a plate reader and CFU counting. The bacterial growth inhibition was observed only in the wells, which were irradiated in presence of biohybrids (Figure 6, A). The CFU counts showed 10^3^ decrease in cells density for the UV-irradiated samples after 18 h of incubation compared to a starting point while a 10^2^ increase was observed for the non-irradiated samples, accounting to a 10^5^ cells density difference among treatments. Moreover, we could also demonstrate efficient inhibition of *E. coli* growth for the biohybrids functionalized with lower concentrations of CIP-photoN_3_ (**6**, 0.01 mM, and 0.005 mM), that fully corresponds to the MIC data and calculated release efficacy (Figure S23-S24, Supporting Information). Furthermore, control experiments provided evidence that neither VAN-photoN_3_ (**3**) or CIP-photoN_3_ (**6**) incubated with algae nor the by-products after UV-irradiation or UV-irradiation itself influence on bacterial growth (Figures S28-S39, Supporting Information). In order to demonstrate the advantage of the system, the experiment using the biohybrids was repeated with an UV-irradiation step during the exponential phase of bacterial growth. Earlier we demonstrated the successful results against *B. subtilis*, this time we expanded the scope to Gram-negative *E. coli*. After incubating the bacteria with biohybrids coated with CIP-photoN_3_ (0.005 mM) for 5 h in 24-wells plates, the system was UV-irradiated at λ = 365 nm for 5 min and the samples were incubated for 13 h more. A clear inhibition of the bacterial growth was observed after the release of the antibiotic for the samples containing the biohybrids compared to the negative control (Figure S40, Supporting Information).

**Figure 6.**
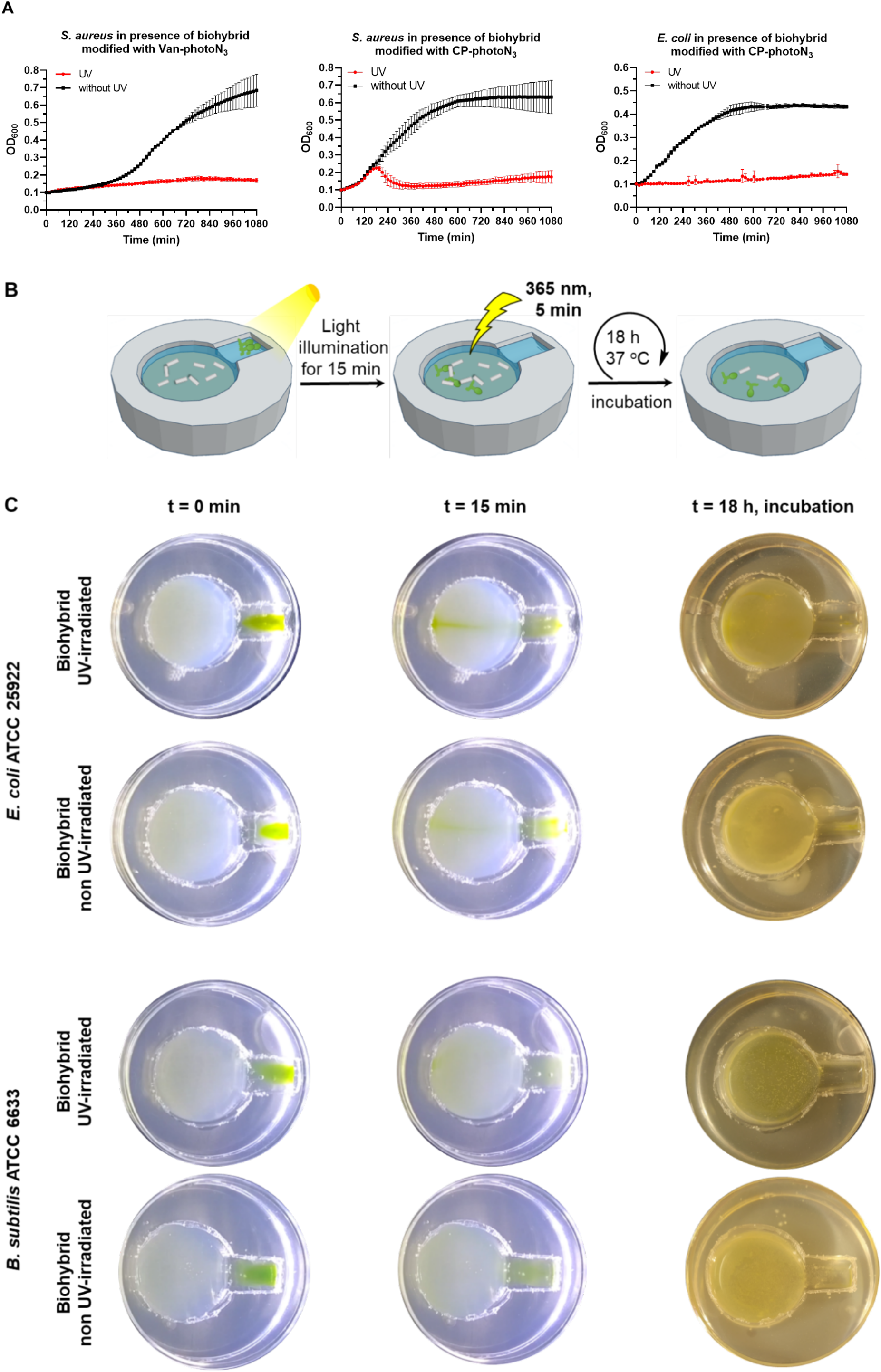
Antibacterial activity of biohybrid *C. reinhardtii*. A) *S. aureus* and *E. coli* growth curves in the presence of biohybrids functionalized with DBCO (0.2 mM) and either VAN-photoN_3_ or CIP-photoN_3_ (0.2 mM). Two samples of functionalized biohybrids were combined when tested against *S. aureus*. The samples were UV-irradiated (λ = 365 nm) at time 0 min for 5 min - red line; the samples were not exposed to a UV-irradiation – black line. Data points adjusted to initial value 0.1 and represent mean value ±SD (n=3). B) Schematics of experiment *in situ*. C) Images of well plate at different time points for the experiment *in situ*.

Finally, the drug delivery experiment by biohybrids was carried out in a special constructed device. A channel connected to a reservoir was built using a Silicone Elastomer Base. The reservoir was filled with MHB agar and the bacteria (*B. subtilis* or *E. coli*, 0.100 μl, OD_600_ = 0.1). The reservoir and the channel were filled with sterile water and the biohybrids were placed at the beginning of the channel. The right side of the channel was illuminated for 15 min to steer biohybrids to the reservoir (Movie S7-S10, Supporting Information). Non-moved algae were removed from the channel and the system was UV-irradiated at λ = 365 nm for 5 min. Then, the cells were incubated at 37 °C for 18 h (Figure 6, B). As a negative control, the samples were not exposed to UV-light. Pleasingly, the bacterial growth inhibition was observed only for the UV-irradiated samples (Figure 6, C). To establish the concept of drug delivery by biohybrids *via* phototaxis, a control experiment was carried out. The biohybrids were placed at the right side of channel in the darkness for 15 min. Only diffusion of the cells inside of the channel was observed. After the removal of the microalgae from the channel, the samples were UV-irradiated and incubated for 18 h. Only the normal bacterial growth was observed, establishing that biohybrid microalgae can be manipulated with external stimulus to deliver their medical cargo with high accuracy (Figure S41-42, Supporting Information). It is well known that UV-light has reduced penetration depth^[71]^ what limits it range on medical application. However, our system could be applicable for the treatment of skin or would infections, as both UV-light and microalgae have been shown to be beneficial and effective against several skin diseases and wound healing.^[72–74]^ Additionally, the change of the photolinker response to near-infrared (NIR) light is a potential solution for a realistic medical application of the designed biohybrids.^[75]^ Moreover, to overcome the issue with low release efficacy of antibiotics by UV-irradiation, other stimuli-responsive release mechanism, such as thermal or chemical triggers may be considered as a future work. Nevertheless, one of the advantages of our system is covalent functionalization of microalgae surface. This approach demonstrated high labeling efficacy and prevented the undesired loss of the cargo during its delivery to the target, which was previously observed for biohybrids modified by non-covalent interactions.^[27,60]^ Moreover, our system can still be potentially functionalized with magnetic nanoparticles to extrinsically manipulate the organism by magnetic control. The attachment can be conducted via electrostatic interactions between negatively charged *C. reinhardtii* cell wall and positively charged magnetic microbeads.^[61]^ Collectively, the data presented in this study provides strong evidence for the directed delivery of antibiotics by designed biohybrid microswimmers stimulated by light.

## Conclusion

In this work, we engineered a new biohybrid microswimmer featuring antibacterial properties. This designed strategy included an operationally simple procedure with high manufacturing yields. The surface functionalization did not affect the microalgae motility retaining phototactic behavior. We demonstrated the application of our biohybrid microswimmers for caging and controlled release of different types of antibiotics, that can expand their use against various bacterial infections. The microswimmers could be guided to a particular region and release their antibacterial cargo with high precision. In summary, these results represent the first reports where microalgae-based biohybrids were used for the guided drug delivery of antibiotics with controlled drug release at bacterial target site.

## Experimental Section

### Chemistry

#### General Methods

Reactions were carried out under inert gas (N_2_ or Ar) in oven-dried (110 °C) glass equipment and monitored for completion by TLC or UHPLC-MS (ESI). Solvents for reactions and analyses were of analytical grade. ciprofloxacin, 5-Hydroxy-2-nitrobenzaldehyde, and 1-azido-2-(2-(2-(2-bromoethoxy)ethoxy)ethoxy)ethane were purchased from Biochemika, FluoroChem, and BroadPharm, respectively. The synthesis of compounds **1**, **3** and **4** have already been reported.^[62,76,77]^ Analytical thin-layer chromatography (TLC) was run on Merck TLC plates silica gel 60 F_254_ on aluminium sheets with the indicated solvent system; the spots were visualized by UV light (λ = 365 nm) and stained by KMnO_4_. Silica gel column chromatography was performed using silica gel 60Å (230-400 mesh) purchased from Sigma-Aldrich with the solvent mixture indicated. The SPE columns used were DSC-18 (Supelco, Sigma). Semi-preparative HPLC separations were performed on a Shimadzu HPLC system (LC-20AP dual pump, CBM-20A 389 Communication Bus Module, SPP-20, A UV/vis Detector, FRC-10A Fraction collector) using reverse-phase (RP) column Synergi Hydro-RP (150 mm × 10 mm; 5 μm, 80 Å). Ultrahigh-performance liquid chromatography (UHPLC) coupled to mass spectrometer (MS) experiments were performed on an Ultimate 3000 LC system (HPG-3400 RS 397 pump, WPS-3000 TRS autosampler, TCC-3000 RS column oven, Vanquish DAD detector from Thermo Scientific) coupled to a triple quadrupole (TSQ Quantum Ultra from Thermo Scientific). The separation was performed using a RP column (Kinetex EVO C18; 50 × 2.1 mm; 1.7 μm; 100 Å, Phenomenex), a flow of 0.4 mL/min, a solvent system composed of A (H_2_O + 0.1% HCO_2_H) and B (MeCN + 0.1% HCO_2_H) and an elution gradient starting with 5% B, increasing from 5-95% B in 3.5 min, from 95-100% B in 0.05 min, and washing the column with 100% B for 1.25 min. In UHPLC-MS measurements after the photolysis experiments, the fragments ions were monitored by SIM mode focusing on the m/z 332.1 and 684.2 Da. Lyophilization was performed on a Christ Freeze-dryer ALPHA 1-4 LD plus. High-resolution electrospray mass spectra (HR-ESI-MS) were recorded on a times TOF Pro TIMS-QTOF-MS instrument (Bruker Daltonics GmbH, Bremen, Germany). The samples were dissolved in (e.g., MeOH) at a concentration of ca. 50 416 μg/mL and analyzed via continuous flow injection (2 μL/ 417 min). The mass spectrometer was operated in the positive (or negative) electrospray ionization mode at 4000 V (−4000 V) capillary voltage and −500 V (500 V) end plate offset with a N_2_ nebulizer pressure of 0.4 bar and a dry gas flow of 4 L/min at 180 °C. Mass spectra were acquired in a mass range from m/z 50 to 2000 at ca. 20 000 resolution (m/z) and at 1.0 Hz rate. The mass analyzer was calibrated between m/z 118 and 2721 using an Agilent ESI-L low concentration tuning mix solution (Agilent, USA) at a resolution of 20 000 giving a mass accuracy below 2 ppm. All solvents used were purchased in best LC-MS quality. 1H and 13C NMR spectra were recorded on Avance II or III-500 (500 MHz with Cryo-BBO, TXI, BBI or BBO probes). Chemical shifts are given in parts per million (ppm) on the delta (δ) scale and coupling constants (J) were reported in Hz. Chemical shifts were calibrated according to the used solvents.^[78]^

#### Ethyl 1-cyclopropyl-6-fluoro-7-(4-(5-hydroxy-2-nitrobenzyl)piperazin-1-yl)-4-oxo-1,4-dihydroquinoline-3-carboxylate (*5*)

To the solution of compound **3** (110 mg, 0.306 mmol) in DCM (3.08 mL), aldehyde **4** (91 mg, 0.430 mmol), NaBH(OAc)_3_ (134 mg, 0.614) and two drops of AcOH were added. The reaction was stirred for 3 h at rt. The reaction mixture was diluted with DCM and poured into sat. NaHCO_3_. The water phase was extracted with DCM (3×20 mL). The organic phase was collected, dried over Na_2_SO_4_, and concentrated under reduced pressure. Ether was added into the crude and the formed precipitate was filtered and dried leading to fully protected intermediate as yellowish powder. This powder (35 mg, 0.063 mmol) was dissolved in dioxane (0.9 mL) and cooled down to 0 °C. 4M HCl in dioxane (0.3mL) was added dropwise. The reaction mixture was stirred at rt for 6 h. The formed precipitate was filtered and washed with ether (2×10 mL) giving the desired product as a yellow solid (33 mg, 0.06 mmol, 96%). **^1^H NMR** (500 MHz, DMSO-*d*_6_) δ 8.44 (s, 1H), 8.24 (d, *J* = 8.2 Hz, 1H), 7.80 (d, *J* = 13.1 Hz, 1H), 7.47 (d, *J* = 6.9 Hz, 1H), 7.26 (s, 1H), 7.13 (d, *J* = 7.9 Hz, 1H), 4.21 (q, *J* = 6.9 Hz, 2H), 4.04 – 3.70 (m, 8H), 3.68 – 3.56 (m, 3H), 3.55 – 3.47 (m, 2H), 3.46 – 3.38 (m, 2H), 1.27 (t, *J* = 6.9 Hz, 3H). **^13^C NMR** (126 MHz, DMSO-*d_6_*) δ 171.49, 164.34, 163.00, 153.31, 151.35, 148.21, 142.18, 140.11, 137.98, 128.84, 127.66, 122.56, 121.22, 117.11, 111.68, 109.34, 106.70, 59.79, 57.12, 51.43, 46.30, 34.79, 14.27, 7.57. **HRMS (ESI)**: calcd for C_26_H_28_O_6_N_4_F [M+H]^+^, *m/z* = 511.19874, found 511.19915.

#### Ciprofloxacin derivative (*8*)

Compound **5** (33 mg, 0.06 mmol) was dissolved in DMF (0.5 mL).

K_2_CO_3_ (17.9 mg, 0.129 mmol) and bromo-PEG_3_-azide (20 mg, 0.07 mmol) were added. The reaction mixture was stirred at 80 °C overnight. The solvent was evaporated and the purification carried out by column chromatography (DCM:MeOH 10:1). The desired product was isolated as a yellowish oil (24 mg, 0.06 mmol, 52%). **^1^H NMR** (500 MHz, MeOD) δ 8.61 (s, 1H), 7.96 (d, *J* = 9.0 Hz, 1H), 7.83 (d, *J* = 13.6 Hz, 1H), 7.46 (d, *J* = 7.3 Hz, 1H), 7.27 (d, *J* = 2.8 Hz, 1H), 7.01 (dd, *J* = 9.0, 2.8 Hz, 1H), 4.32 (q, *J* = 7.1 Hz, 2H), 4.27 – 4.24 (m, 2H), 3.92 (s, 2H), 3.90 – 3.87 (m, 2H), 3.73 – 3.69 (m, 3H), 3.69 – 3.58 (m, 13H), 3.36 – 3.32 (m, 2H), 2.73 – 2.67 (m, 4H), 1.37 (t, *J* = 7.1 Hz, 3H), 1.17 – 1.13 (m, 2H). **^13^C NMR** (126 MHz, MeOD) δ 175.43, 166.22, 163.67, 153.89, 149.87, 146.42, 146.33, 144.15, 139.87, 137.61, 128.42, 118.02, 114.19, 113.09, 112.90, 110.41, 106.98, 71.81, 71.67, 71.55, 71.14, 70.58, 69.36, 61.69, 60.34, 54.09, 51.77, 36.25, 14.69, 8.54. **HRMS (ESI)**: calcd for C_34_H_43_O_9_N_7_F [M+H]^+^, *m/z* = 712.31008, found 712.30991.

#### Ciprofloxacin derivative (*6*)

Compound **8** (8 mg, 0.011 mmol) was dissolved in methanol (0.090 mL) and water (0.030 mL). LiOH (2.7 mg, 0.112 mmol) was added, and the reaction mixture was stirred at 80 °C for 2 h. The reaction was acidified by addition of 1M HCl. The methanol was evaporated, and the rest was extracted with DCM (3×10 mL). The organic phase was dried over Na_2_SO_4_ and concentrated under reduced pressure. The crude was dissolved in mixture of water/CH_3_CN+0.1%FA filtered through syringe filter (1.0 μm) and purified by semi-prep RF-HPLC: gradient 10% B for 7 min; 10% - 45% B for 28 min; 45% - 100% B for 5 min, wash. The desired product, eluting at 32.2 min, was collected and lyophilized to afford product (3.2 mg, 0.011 mmol, 42%) as a yellowish oil. **^1^H NMR** (500 MHz, DMSO-*d*_6_) δ 15.21 (s, 1H), 8.65 (s, 1H), 7.98 (d, *J* = 9.0 Hz, 1H), 7.89 (d, *J* = 13.6 Hz, 1H), 7.56 (d, *J* = 7.3 Hz, 1H), 7.27 (d, *J* = 2.8 Hz, 1H), 7.06 (dd, *J* = 9.0, 2.8 Hz, 1H), 4.28 – 4.18 (m, 2H), 3.88 (s, 2H), 3.84 – 3.74 (m, 3H), 3.64 – 3.48 (m, 12H), 2.65 – 2.61 (m, 4H), 1.31 (t, *J* = 6.5 Hz, 2H), 1.19 – 1.12 (m, 2H). **^13^C NMR** (126 MHz, DMSO) δ 176.36, 165.96, 161.86, 154.04, 152.06, 148.00, 145.20, 142.14, 139.18, 136.20, 127.41, 118.71, 116.69, 113.10, 111.00, 106.50, 69.94, 69.84, 69.81, 69.69, 69.25, 68.68, 68.01, 58.39, 52.40, 49.99, 49.52, 49.49, 35.88, 7.57. **HRMS (ESI)**: calcd for C_32_H_39_O_9_N_7_F [M+H]^+^, *m/z* = 684.27878, found 684.27904.

### Biology

#### Cultures and growth conditions

*Chlamydomonas reinhardtii* (*C*. *reinhardtii*, 11-32b) was purchased from EPSAG (Experimentelle Phykologie und Sammlung von Algenkulturen der Universität Göttingen) and grown in TAP medium (50 mL) prepared according to the published procedure (2.42 g Tris, 25 mL salts stock, 0.375 mL Phosphate Solution, 1.0 mL Hutner’s tarce metals, 1.0 mL glacial AcOH).^[79]^ The algae were grown at 25 °C under constant illumination of 8 μmol photons/m^2^s at 150 rpm on a rotary shaker. *Bacillus subtilis* (*B. subtilis*, ATCC 6633), *Escherichia coli* (*E. coli*, ATCC 25922), *Staphylococcus aureus* (*S. aureus*, ATCC 43300, MRSA) was purchased from the German Collection of Microorganisms and Cell Cultures (DSMZ) or the American Type Culture Collection (ATCC). The bacteria culture was stored at –80 °C, and new cultures were prepared by streaking on Mueller Hinton Agar plates. The overnight culture was prepared by inoculating a single colony into a plastic sterile tube (5 mL) containing the LB medium for *E. coli* and MHB medium for *B. subtilis* and *S. Aureus*. The cultures were shaken (200 rpm/min) overnight at 37 °C.

#### Preparation of the biohybrids

The algae were grown for 4 days and the cell surface modification was carried out in two steps. The washing steps was carried out with dTAP (TAP:H_2_O = 1:10). Non-modified algae for the control experiments were exposed to the similar functionalization steps but excluded the addition of any compounds. *1^st^ functionalization step:* An microalgae culture (0.5 mL) with a cell density of 1.5×10^6^ cells/ml calculated using a counting chamber was centrifuged (5 min, 4000 rcf/min) and the supernatant was removed. The algae cells were washed with dTAP (1 mL), centrifuged (5 min, 4000 rcf/min) and the supernatant was removed, this step was repeated one more time. The cells were resuspended in a NHS-PEG_4_-dibenzocyclooctyne (DBCO) solution (dTAP, 0.250 mL) at the concentrations 10 times higher than antibiotic. The mixture was shaken for 1h, the samples were centrifuged (3 min, 2000 rcf/min) and the supernatant was removed. The pellet was washed with dTAP (1 mL), centrifuged (3 min, 2000 rcf/min) and the supernatant was removed, this step was repeated 2 times. *2^d^ functionalization step*: The modified algae were resuspended in a solution of either a vancomycin azide derivative (dTAP, 0.250 mL, with 0.002% Tween 80) or ciprofloxacin derivative (dTAP, 0.5 mL) at the desired concentrations. The mixture was incubated for 1 h, the samples were centrifuged (3 min, 2000 rcf/min), the supernatant was removed, the samples were washed (dTAP, 1 mL), centrifuged (3 min, 2000 rcf/min) and the supernatant was removed. The washing step was repeated 6 times.

#### Photolysis Experiment

Photolysis experiments were performed using a Sina UV lamp (SI-MA-032-W; equipped with UV lamps 4 × 9, 365 nm) at the distance of ~5 cm. See the Supporting Information for the detailed procedure.

#### Motility quantification experiment

The wild type and biohybrid microalgae were preserved in water (1 mL) for 1 h. Then, the cells were centrifuged (3 min, 2000 rcf/min) and water was removed. The cells were resuspended in fresh water (0.500 mL). In glass slide, the frame-seal slide chamber (15×15 mm) was attached. The wild type algae or biohybrids (65 μl) was added, and the chamber was sealed with a cover slip. The 2D swimming speed analysis and swimming trajectories of wild type and biohybrid microalgae were recorded by the Olympus IXplore Spin SR10 spinning disk confocal super resolution microscopy (Lase line: 488 (100 mW) 8% power; objective UPLSAPO UPlan S Apo 30×). The sequential images were recorded in a fixed interval of 100 ms per frame. The 2D swimming trajectories and the calculation of speed were done by TrackMate FIJI.

#### Negative phototaxis studies

The modified microalgae (1 mL) or control algae (1 mL) were placed into the cell culture wells (r = 25 mm). The wells were separated in the middle and the algae were collected from the right and the left side of the well. The cell density (OD_750_) was analyzed by a plate reader (Synergy H1 apparatus from BioTek). The cells were placed back into the cell culture well and the right side of the well was illuminated for 5 min to steer microalgae to the left. The wells were split again in the middle and the algae were collected from both sides to analyze the difference in OD_750_. The microalgae counts were calculated from the calibration curve (Figure S9, Supporting Information).

#### MICs of Tested Compounds

The minimum inhibitory concentration (MIC) of tested compounds was determined using the broth microdilution method according to the guidelines outlined by the European Committee on Antimicrobial Susceptibility Testing (EUCAST).^[80]^ See the Supporting Information for the detailed procedure.

#### Antibacterial activity of biohybrid microalgae. General protocol

The overnight bacterial culture was diluted to an optical density (600 nm) of 0.1 by adding a mixture of biohybrid microalgae in TAP (0.250 ml) and MHB (0.250 ml). The mixture was UV-irradiated at λ=365 nm for 5 min (unless stated otherwise) and grown in a 24-wells plate at 37 °C. The cell density (600 nm) was measured every 15 min for 18 h (with shaking between measurements) in a microplate reader. All experiments were performed in triplicates. For the detailed procedure of the control experiments see the Supporting Information.

Calculation of colony forming units (CFU) was carried out for bacterial culture at time 0 and 18 h (after the experiment with microalgae) by 10-fold dilutions prepared in PBS. 100 μl of suspension was drop-plated on the MHB agar. CFU were enumerated after 24 h of incubation at 37 °C. The experiment was performed in triplicate. The final number represented as an average value from three experiments.

#### Drug delivery experiment

SYLGARD 184 Silicone Elastomer Base was used for the casting process to obtain the desired device. The surface was modified by the oxygen plasma treatment in the plasma reactor for 10 min at 350V. The form was disinfected with ethanol and placed into a Petry dish (r = 50 mm). The reservoir was filled with MHB agar (6 mL) followed by the addition of the bacterial suspension (*E. coli* ATCC 25922, *B. subtilis* ATCC 6633, 1×10^8^ cfu/mL, 0.100 μL). The biohybrid microalgae (50 μL) was carefully placed on the right side of the channel. The light stimulus was switched to the right for 15 min. The movement was recorded in time-lapse mode, and lately accelerated 8 times. The microalgae remained in the channel was carefully removed and the plates were either UV-irradiated for 5 min at λ = 365 nm or was not exposed to UV-light. Next, the plates were incubated for 18 h at 37 °C. The bacterial growth was determined by visual inspection. For the detailed procedure of the control experiments see the Supporting Information.

## Supporting Information

Supporting Information is attached below.

## Author Contributions

I.S.S. and K. G. designed the study. I.S.S. carried out the synthesis of chemicals, the manufacturing of the biohybrid microswimmers, and all biological experiments. J. V. D. M. performed microscopy experiments of the motility, calculated the velocity, and recorded the velocity movies. I. S. S. and K. G. analyzed data and discussed the results. I. S. S. and K. G. wrote the manuscript.

## Acknowledgements

The authors acknowledge the Swiss National Science Foundation (SNSF, Grant No. 182043), a Bundesstipendium (to I. S. S. and J. V. D. M.), and the Gebert Rüf Stiftung (Microbials: project GRS–061/18) for financial support. The authors acknowledge the NMR and Mass spectrometry facilities, the Center for Microscopy and Image Analysis (ZMB) of the University of Zurich for training and maintenance of the instruments. We thank Mikhail Volkov for the generation of the table of content artwork. Esmael Balaghi from the University of Zurich is thanked for the oxygen plasma treatment of the bacterial reaction vessel.

## Biohybrid Microswimmers Against Bacterial Infections

Biohybrid microswimmers are designed and engineered that carry antibiotic cargo against both gram-positive and gram-negative bacteria. Guided by an external beacon, these algal microrobots deliver the antibiotic payload to the site of bacterial infection, with high spatial and temporal precision, released on-demand by an external trigger.

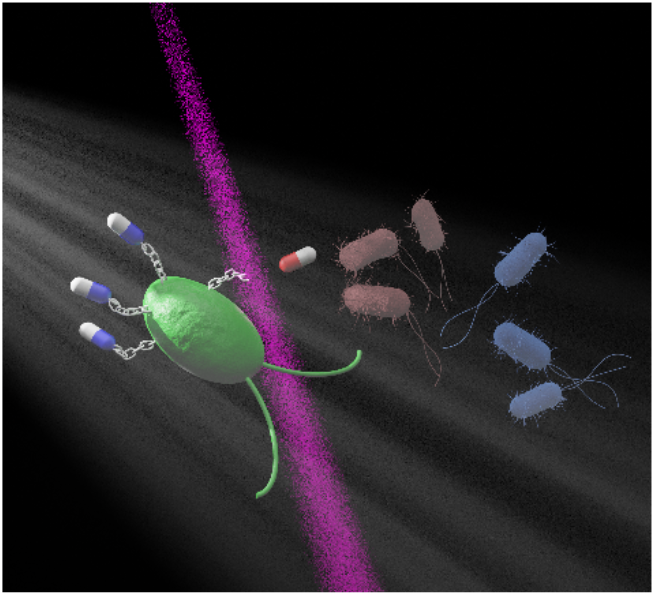

## 1. NMR spectrums of newly synthesised compounds

**Figure S1.**
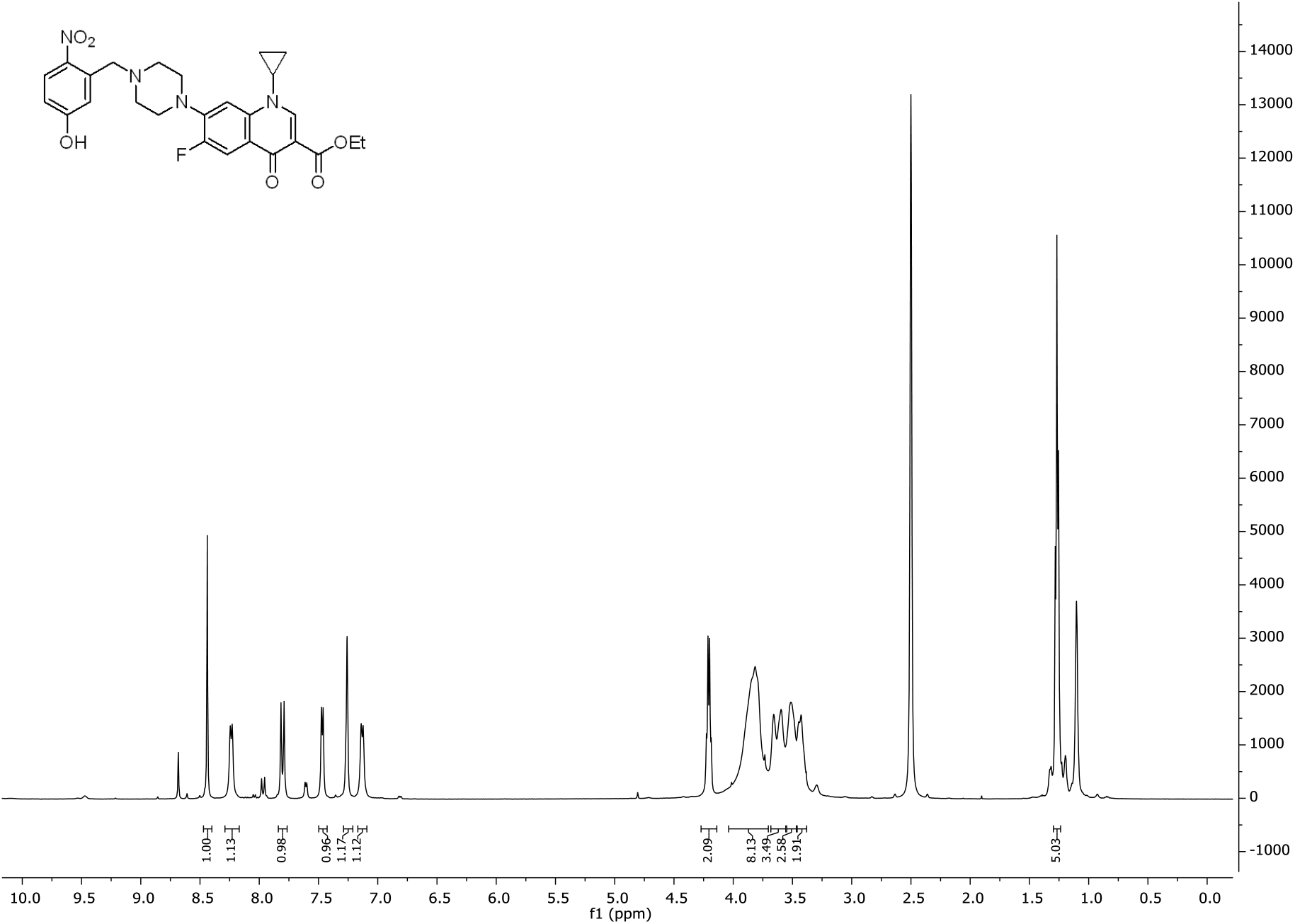
500 MHz ^1^H-NMR spectrum of **5** in DMSO-*d_6_*.

**Figure S2.**
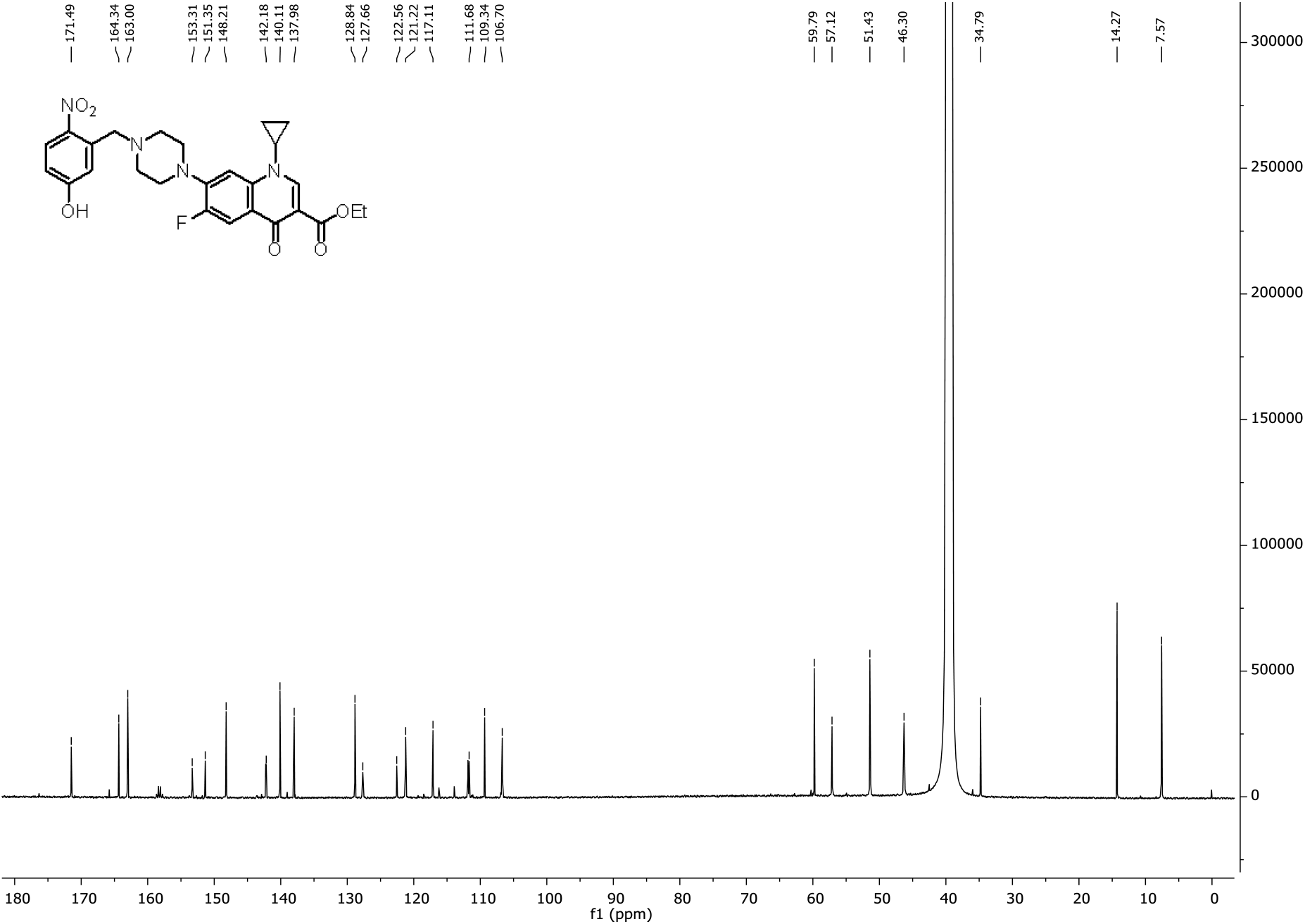
126 MHz ^13^C-NMR spectrum of **5** in DMSO-*d_6_*

**Figure S3.**
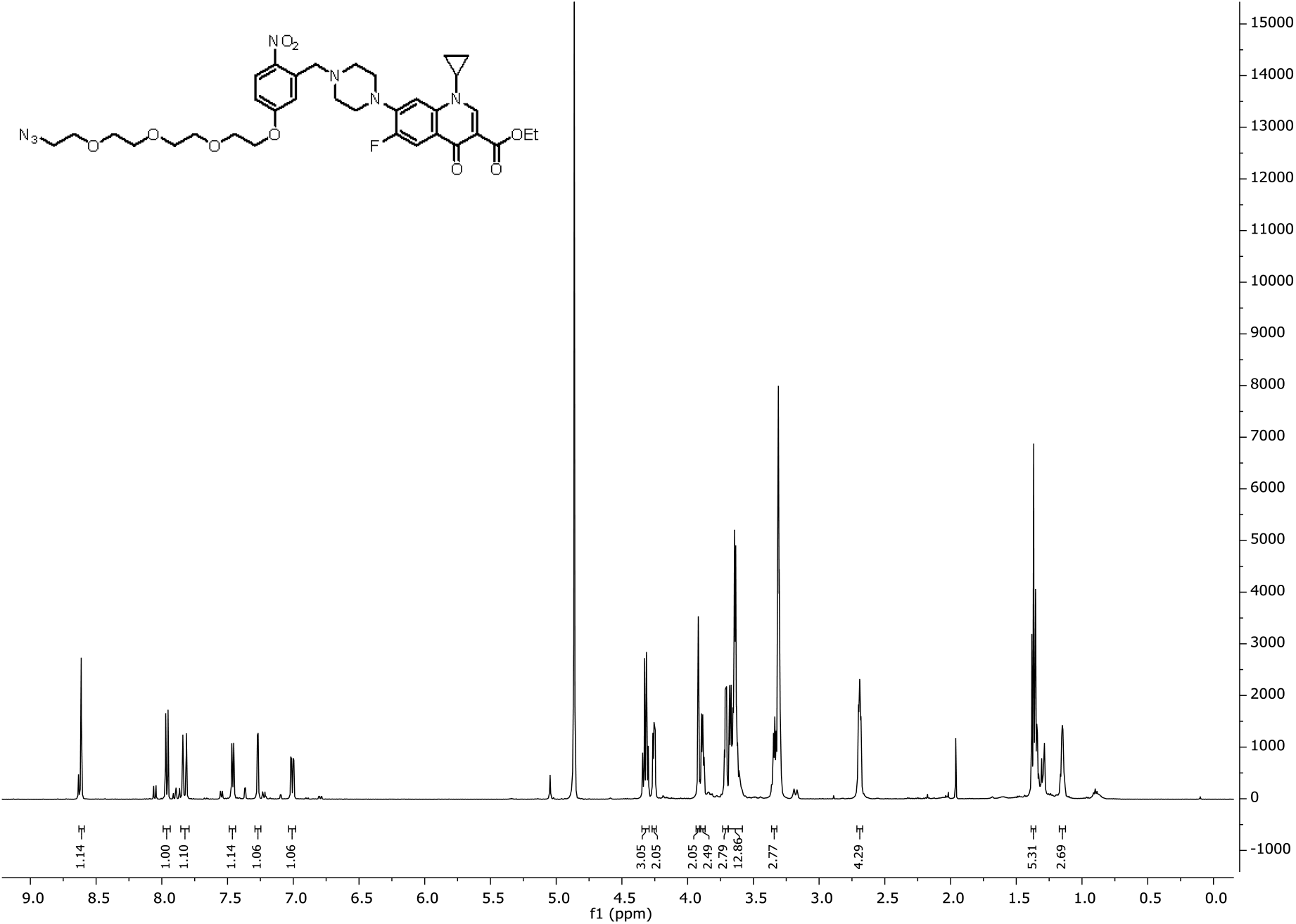
500 MHz ^1^H-NMR spectrum of **8** in MeOD

**Figure S4.**
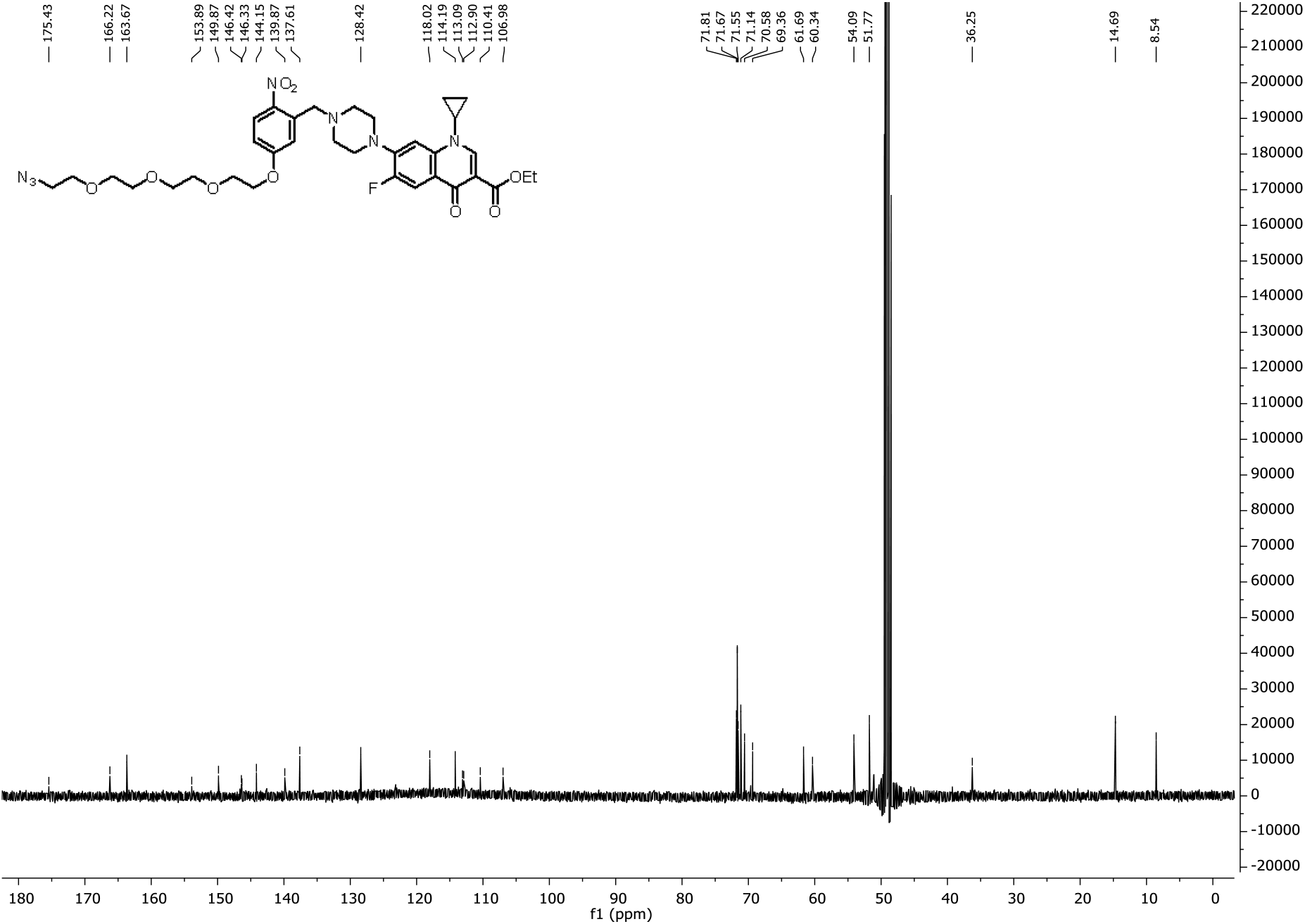
126 MHz ^13^C-NMR spectrum of **8** in MeOD

**Figure S5.**
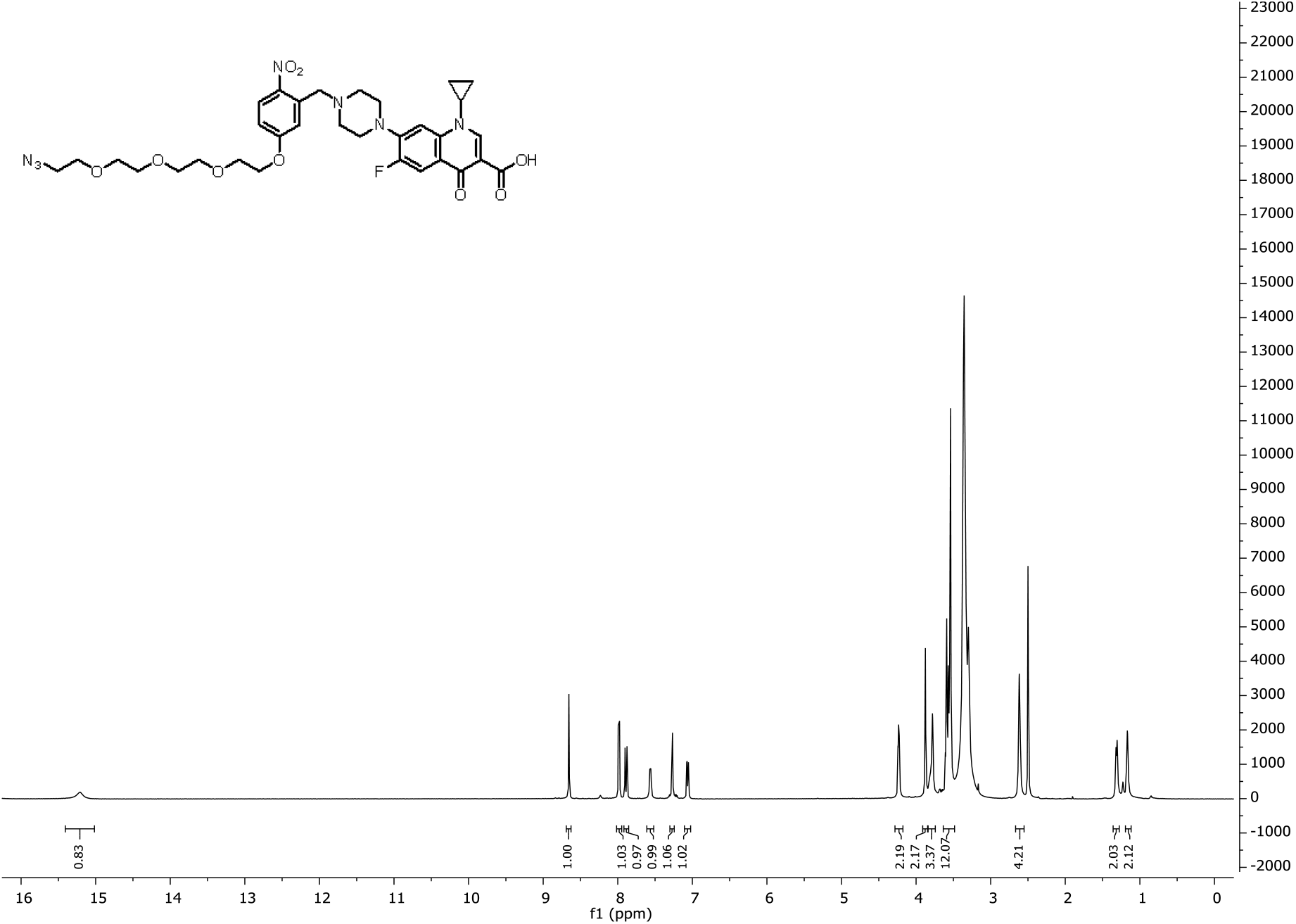
500 MHz ^1^H-NMR spectrum of **6** in DMSO-*d_6_*

**Figure S6.**
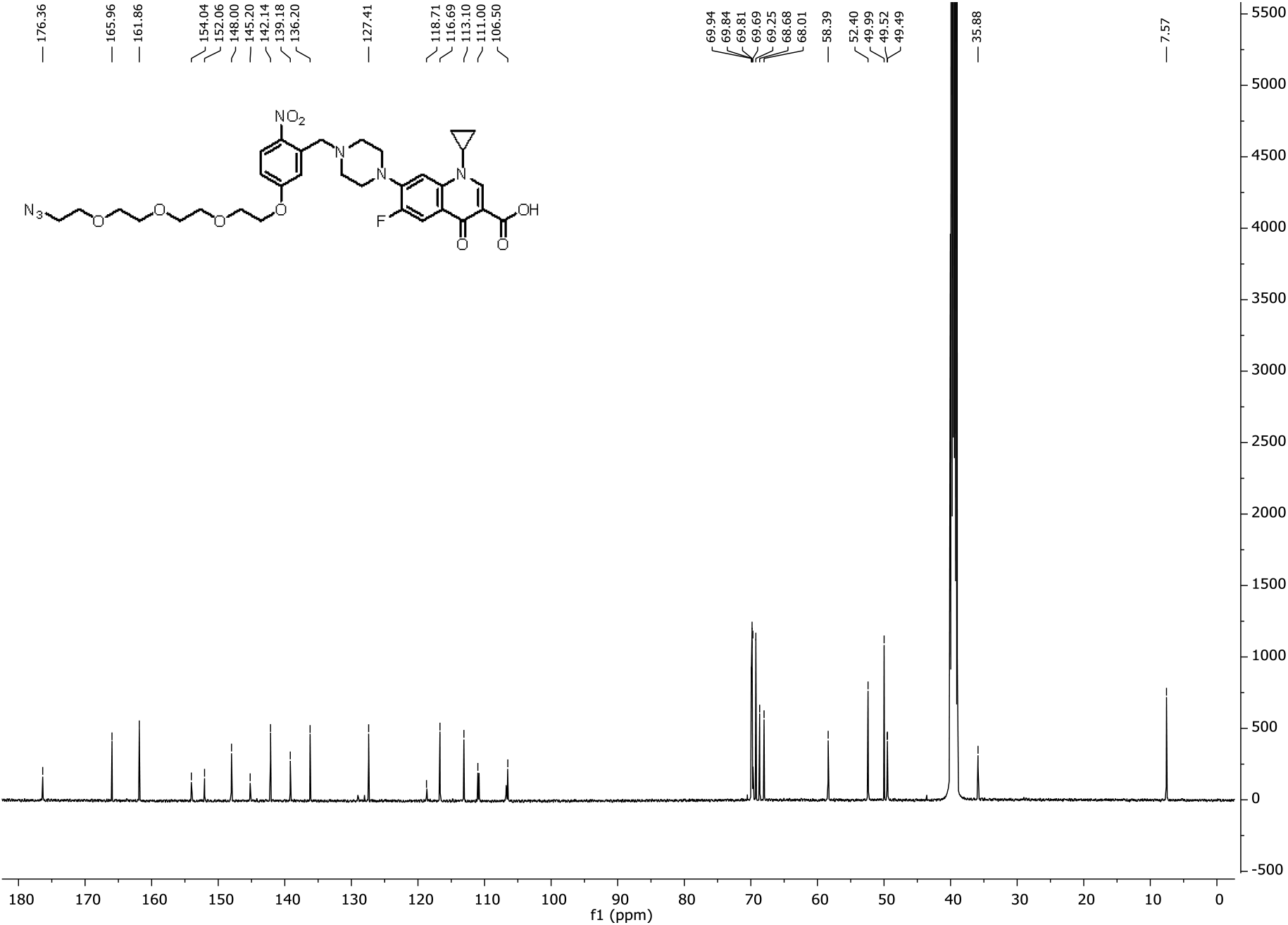
126MHz ^13^C-NMR spectrum of **6** in DMSO-*d_6_*

## 2. Photolysis experiments

The formation of products during the UV-irradiation step of CIP-photoN_3_ the efficiency of photocleavage and the degradation of CIP-photoN_3_ during different time of UV-irradiation were investigated.

**Scheme S7.**
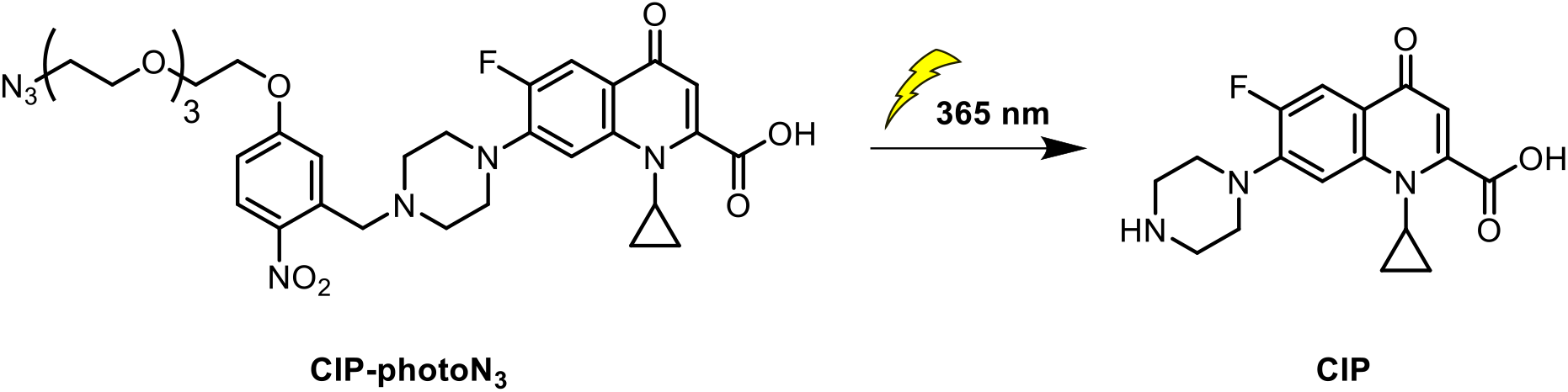
The formation of ciprofloxacin after the UV-irradiation of ciprofloxacin derivative CIP-photoN_3_

**Figure S8.**
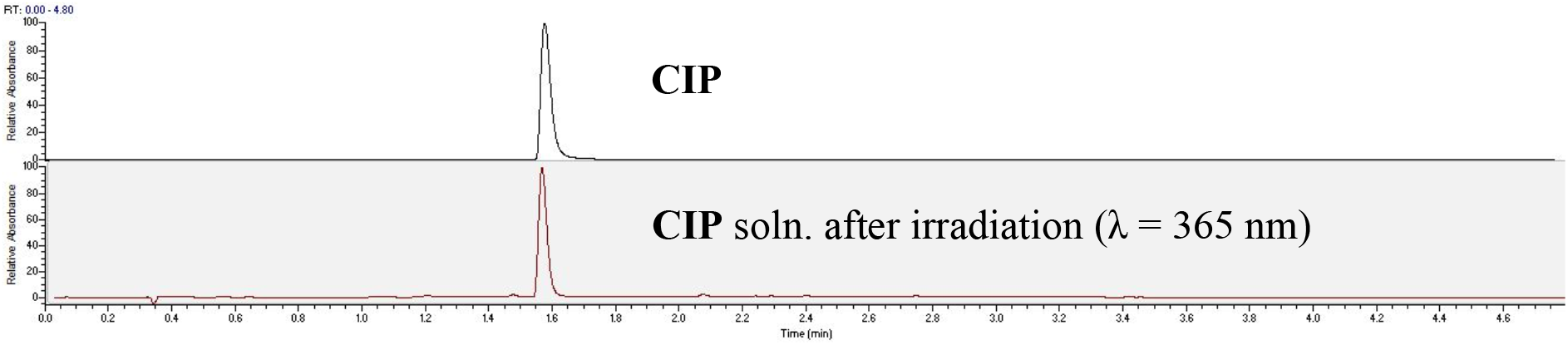
Comparison of UHPLC-MS chromatograms recorded in SIM mode for CIP. From top to bottom: commercially available CIP and the product after UV-irradiation of the CIP-photoN_3_ (**6**) solution.

### 2.1 Quantification protocol of CIP and CIP-photoN_3_

Stock solutions (1 mM) of ciprofloxacin derivative (CIP-photoN_3_ (**6**)) and ciprofloxacin (CIP) were prepared in TAP medium and diluted to obtain several solutions in the concentration range of 0.3 – 10 μM. These solutions were used for the experiments described below and for building the calibration curves. All stock solutions were freshly prepared. Calibration curves were obtained by plotting linear regression of the mass intensity versus the concentration of the standard. From these curves the coefficients of correlation (R^2^) and slope were calculated (Figures S2).

**Figure S9.**
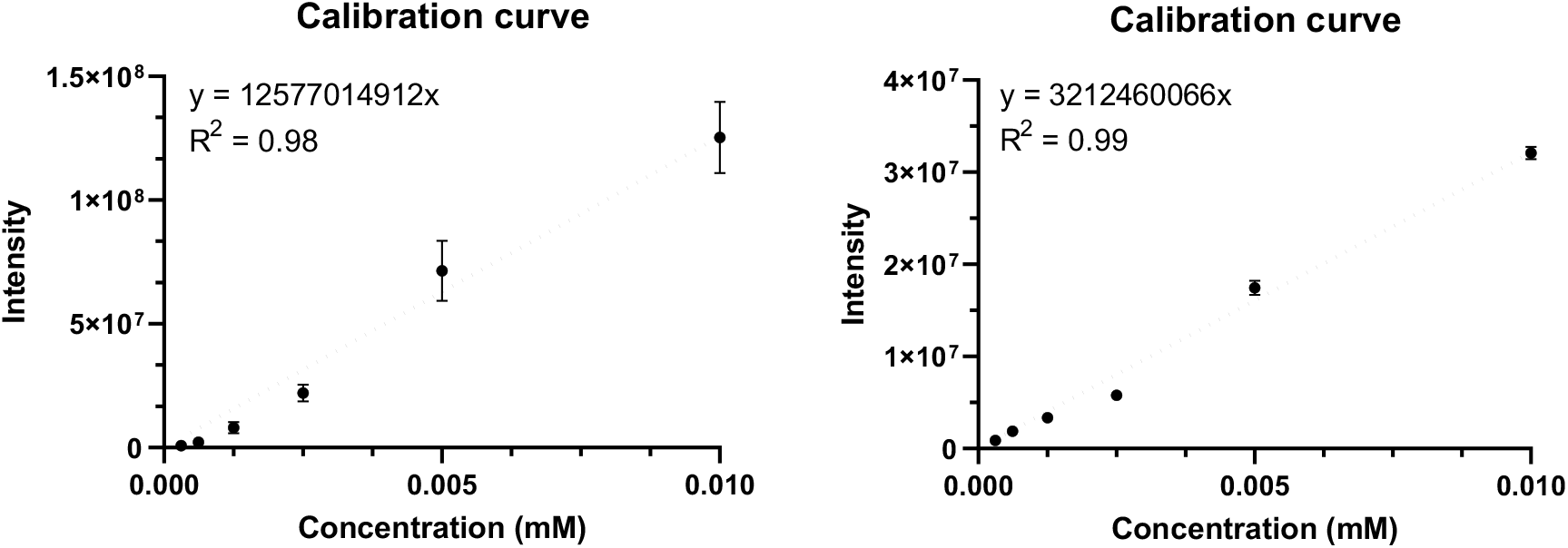
Calibration curves for CIP-photoN_3_ (**6**) on the left and for CIP on the right for the experiments (Section 2.1). Data points represent mean value ± SD (n = 3).

### 2.2 Photolysis experiment of CIP-photoN_3_

CIP-photoN_3_ (**6**) was dissolved in TAP medium (0.5 mL, 10 μM) and loaded in a 24-well plate. The solutions were UV-irradiated at λ = 365 nm. A sample (0.5 mL) was taken after different time of irradiation. The reaction was filtered and analyzed by UHPLC-MS.

**Figure S10.**
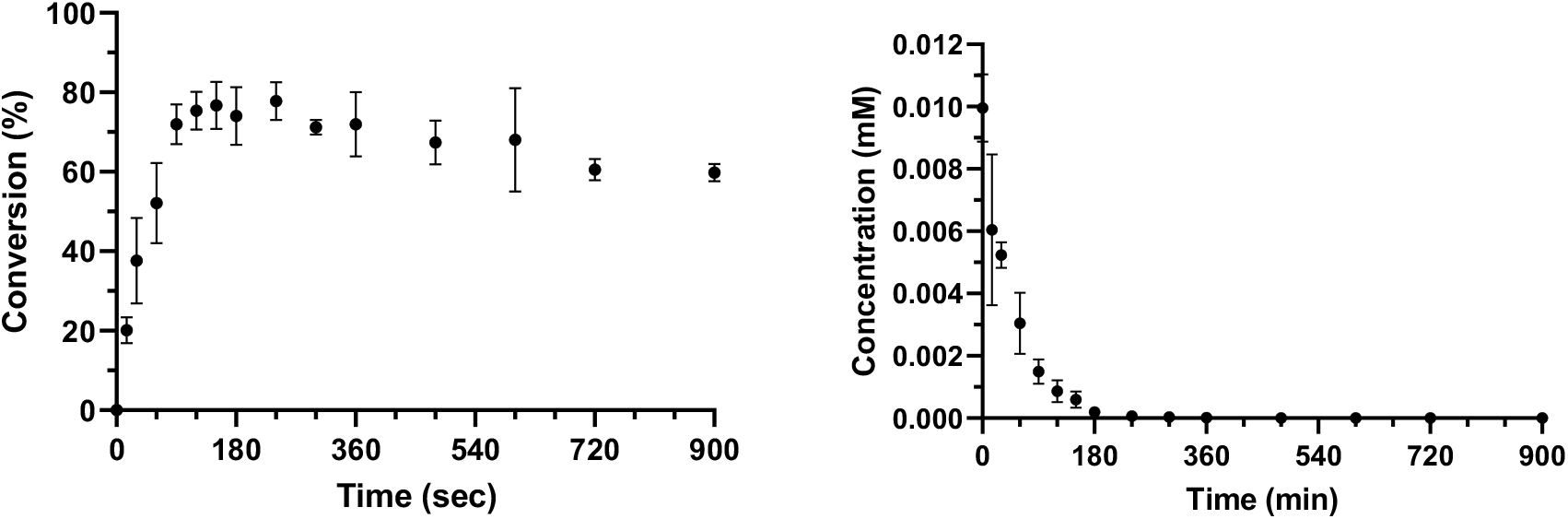
Left: Analysis of CIP-photoN_3_ (**6**) conversion into CIP after the sample irradiation at λ = 365 nm during different time frames. Right: Analysis of CIP-photoN_3_ (**6**) concentration after the sample irradiation at λ = 365 nm during different time. Data points represent mean value ± SD (n = 3).

## 3. Surface modification of *Chlamydomonas reinhardtii*

### 3.1 Screening of the reaction conditions for the algae modification with the ciprofloxacin derivative

*Calculation of labeling efficacy*

The efficacy of surface functionalization was calculated by evaluating the percentage of reacted either VAN-photoN_3_ (**2**) or CIP-photoN_3_ (**6**) after the second step of *C. reinhardtii* surface functionalization. The microalgae were modified by the procedure described in the experimental section in the manuscript, Preparation of the biohybrids section. After the second modification step using either VAN-photoN_3_ (**2**) or CIP-photoN_3_ (**6**), the cells were centrifuged (3 min, 2000 rcf/min), the supernatant was removed and the cells were washed with the diluted media dTAP (1 mL), centrifuged (3 min, 2000 rcf/min) and the supernatant was removed. The washing step was repeated 6 times. The supernatants after the first centrifugation and after each washing step were collected, combined, filtered, and analyzed by UHPLC-MS.

**Table S1:**
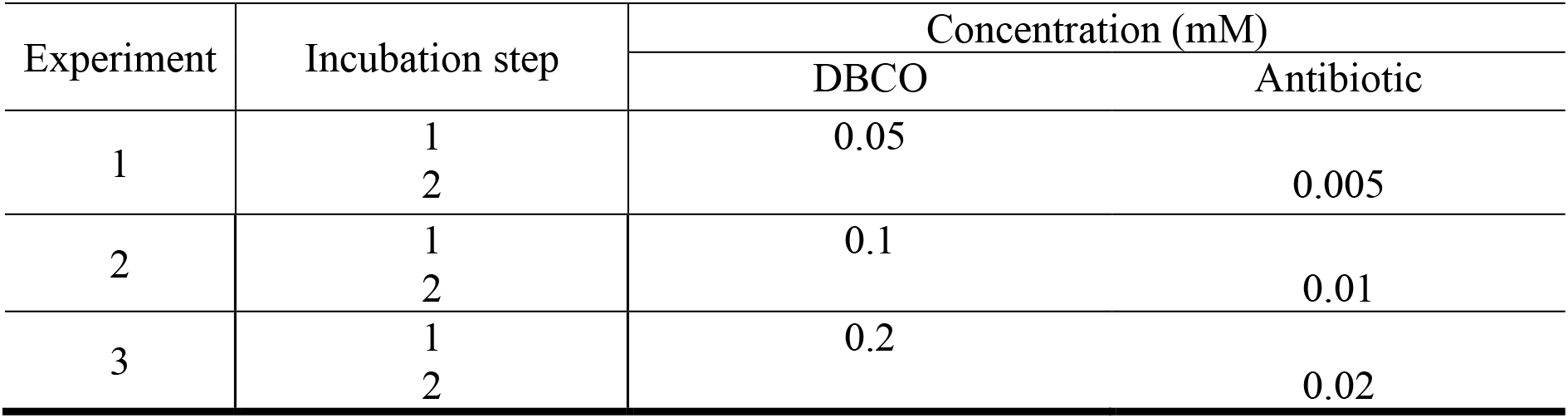
Concentrations of DBCO and synthesized antibiotic derivatives (**2** or **6**) used for *C*. *reinhardtii* surface functionalization

**Figure S11.**
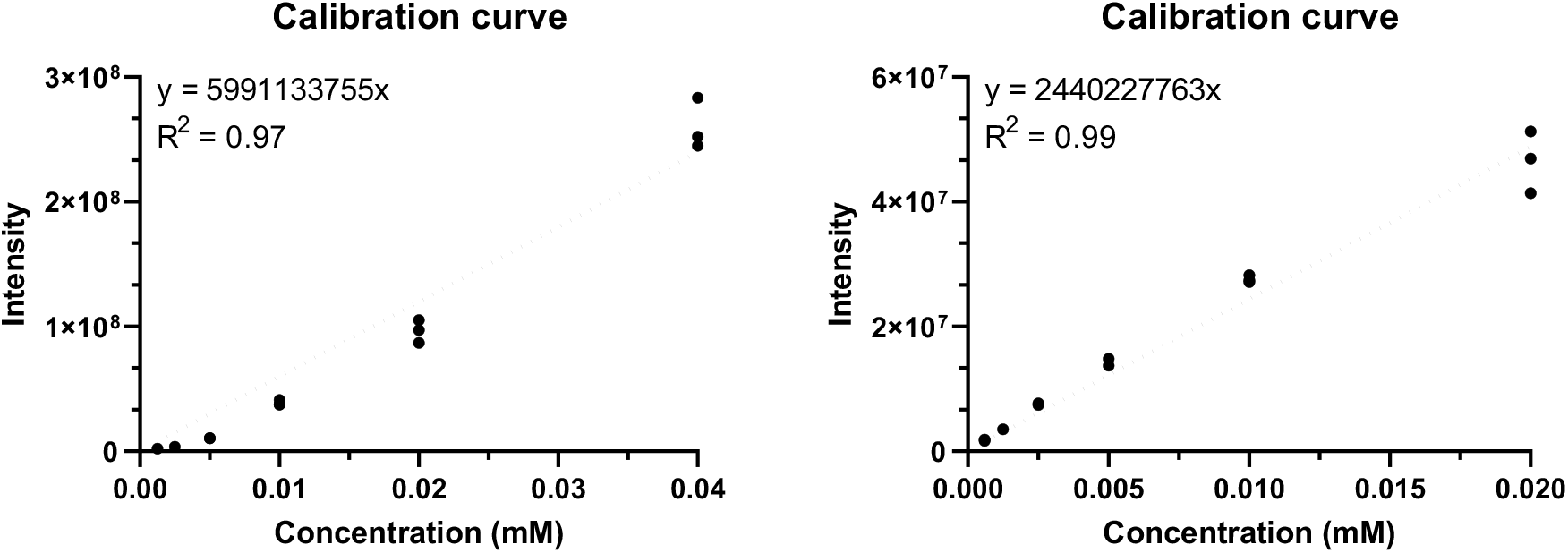
Calibration curves for CIP-photoN_3_ (**6**) on the left and for CIP on the right for the experiments (**Table S1**). Data points represent mean value ± SD (n = 3).

**Figure S12.**
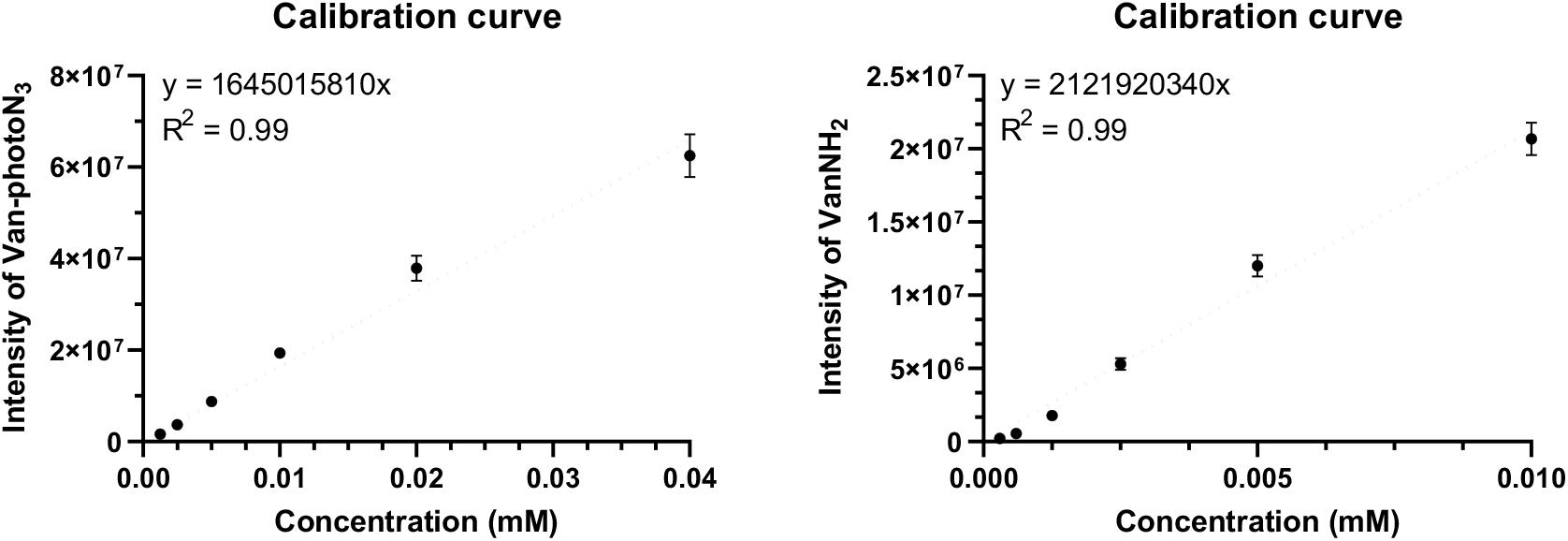
Calibration curves for VAN-photoN_3_ (**2**) on the left and for VAN-NH_2_ (**3**) on the right for the experiments (**Table S1**). Data points represent mean value ± SD (n = 3).

**Figure S13.**
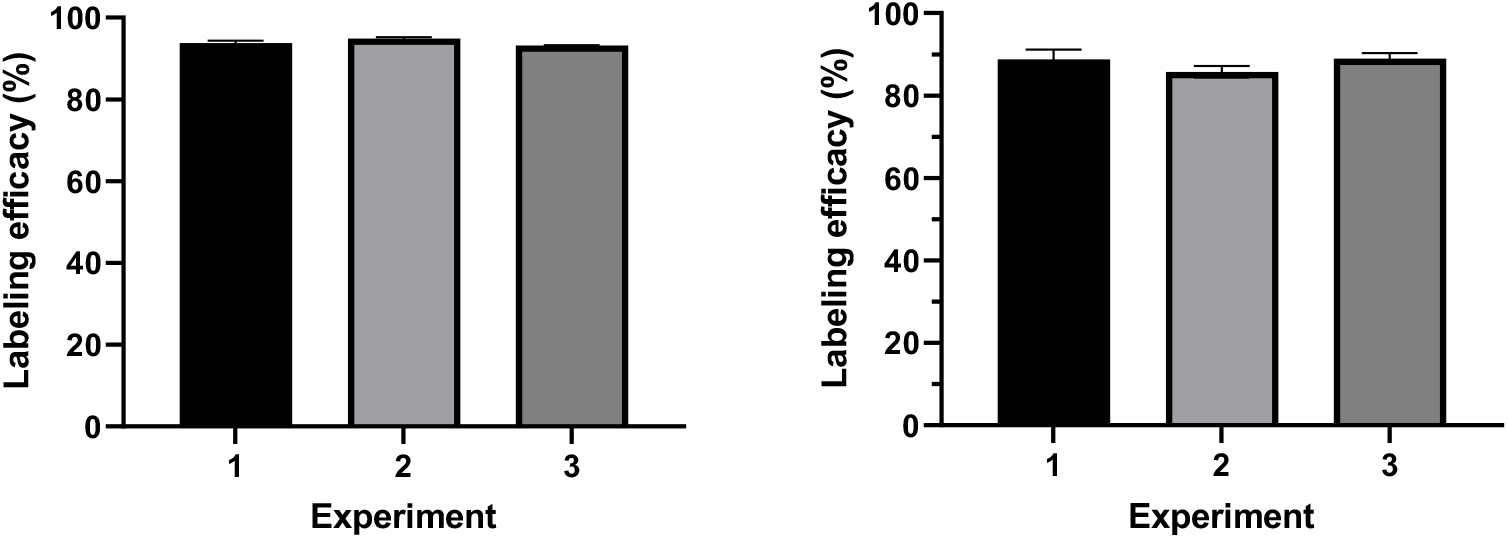
Percentage of microalgae surface labeling efficacy for the experiments (Table S1). Left: Microalgae functionalized with VAN-photoN_3_ (**2**); Right: Microalgae functionalized with CIP-photoN_3_ (**6**). Data are represented mean value ± SD (n = 3).

*Calculation of the release efficacy*

The released concentration of antibiotic from the functionalized surface of *C*. *reinhardtii* was determined by using the calibration curves (Figure S4-S5). The released antibiotic was calculated according to the following equation:

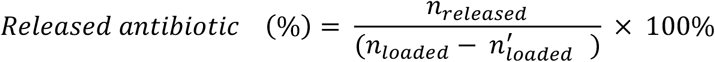

Where *n*_*released*_ is the released amount of either VAN-NH_2_ (**3**) or CIP after irradiation; *n*_*loaded*_ is the amount of either VAN-photoN_3_ (**2**) or CIP-photoN_3_ (**6**) used for the second step of *C*. *reinhardtii* modification; *n*^′^_*loaded*_ is the amount of unreacted either VAN-photoN_3_ (**2**) or CIP-photoN_3_ (**6**) determined by UHPLC analysis (SIM mode).

**Figure S14.**
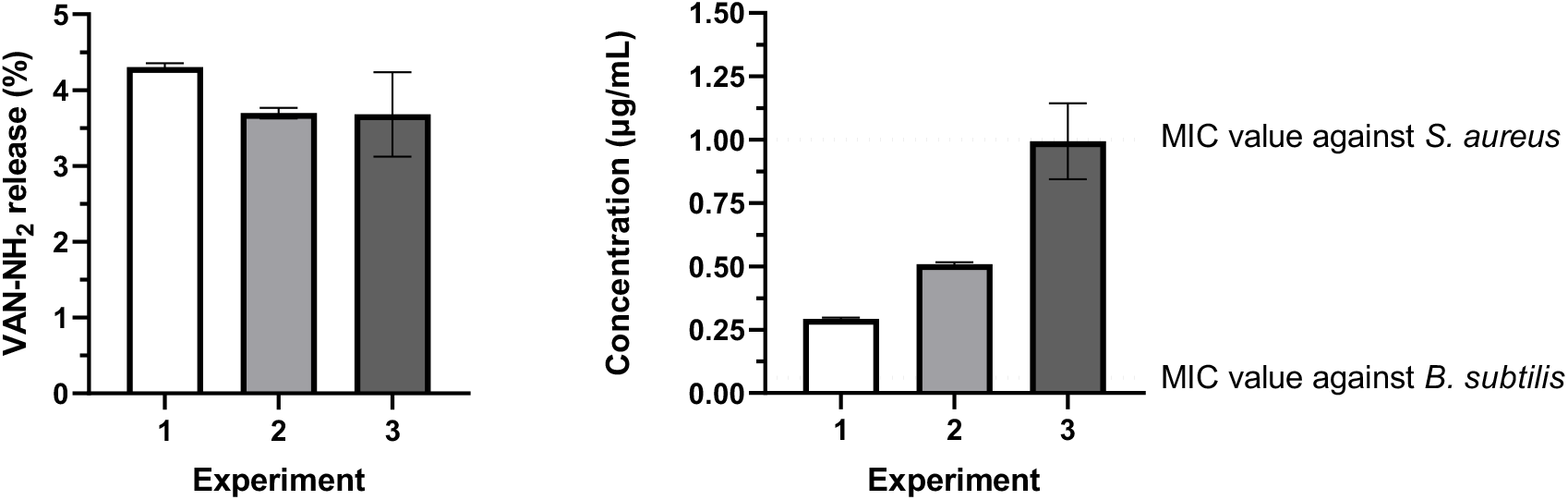
Release of VAN-NH_2_ (**3**) from the microalgae surface after 5 min of UV-irradiation λ = 365 nm for the experiments (**Table S1**). Left: Percentage of VAN-NH_2_ (**3**) released. Right: The concentration of VAN-NH_2_ (**3**) released. The dot line corresponds to MIC values for VAN-NH_2_ (**3**) against *B. subtilis* ATCC 6633 (0.06 μg/mL) and *S. aureus* ATCC 43300 (MRSA) (1 μg/mL). Data are represented mean value ± SD (n = 3).

**Figure S15.**
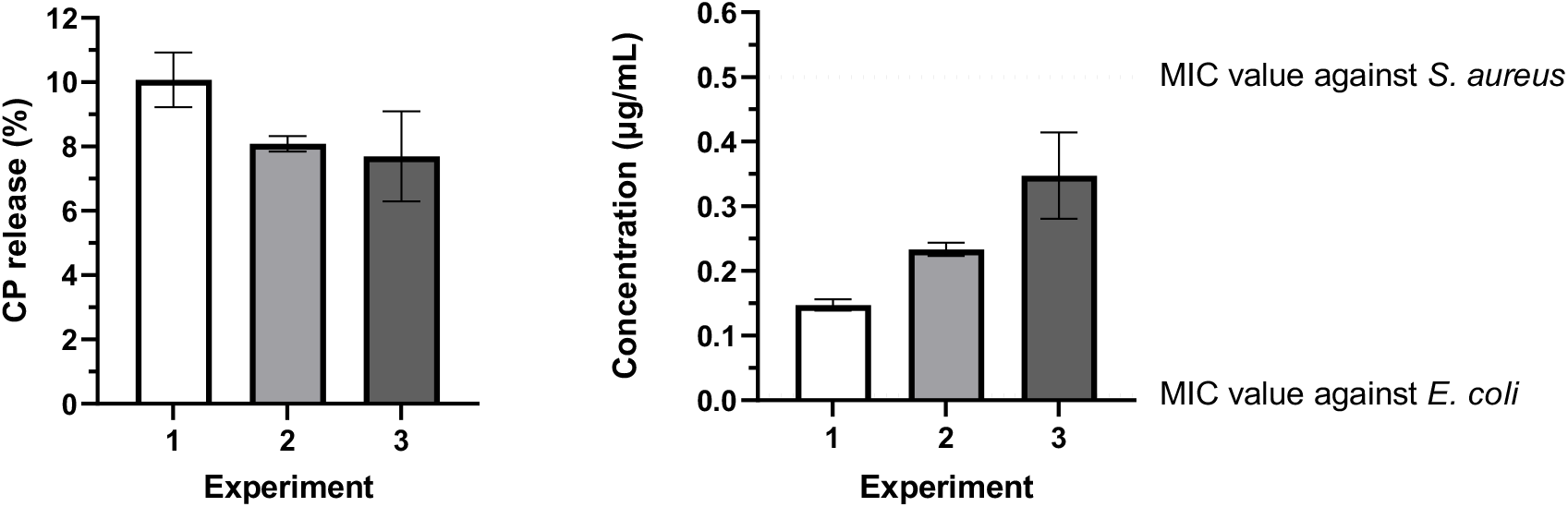
Release of CIP from the microalgae surface after 5 min of UV-irradiation λ=365 nm for the experiments (Table S1). Left: Percentage of CIP released. Right: The concentration of CIP released. The dot line corresponds to MIC values for CIP against *E. coli* ATCC 25922 (0.008 μg/mL) and *S. aureus* ATCC 43300 (MRSA) (0.25-0.5 μg/mL). Data are represented mean value ± SD (n = 3).

## 4. Characterization of biohybrid *Chlamydomonas reinhardtii*

### 4.1 Activation of microalgae motility

The microalgae after the functionalization steps were resuspended in water (1 mL) and preserved for 1 h. Then the cells were centrifuged (3 min, 2000 rcf/min) and the water was removed. The cells were resuspended in the amount of water required for the movement experiment.

### 4.2 Negative phototaxis studies of biohybrid C. reinhardtii

**Figure S16.**
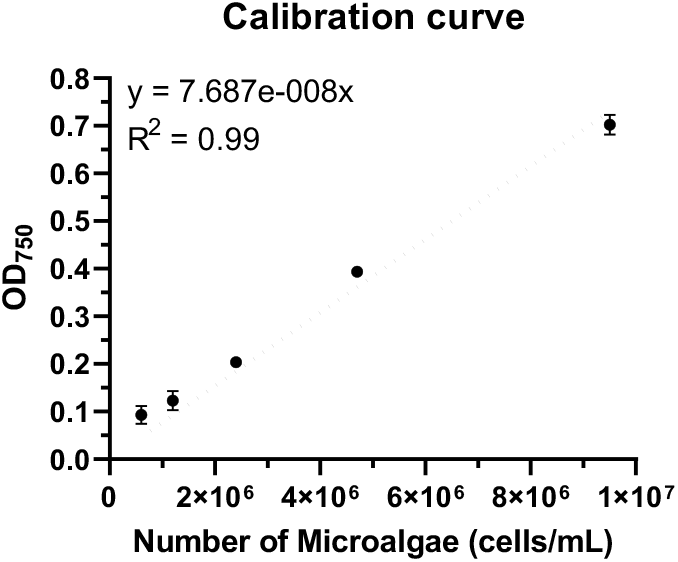
Calibration curve for the calculation of number of algae from optical density data. Data points represent mean value ± SD (n = 3).

**Figure S17.**
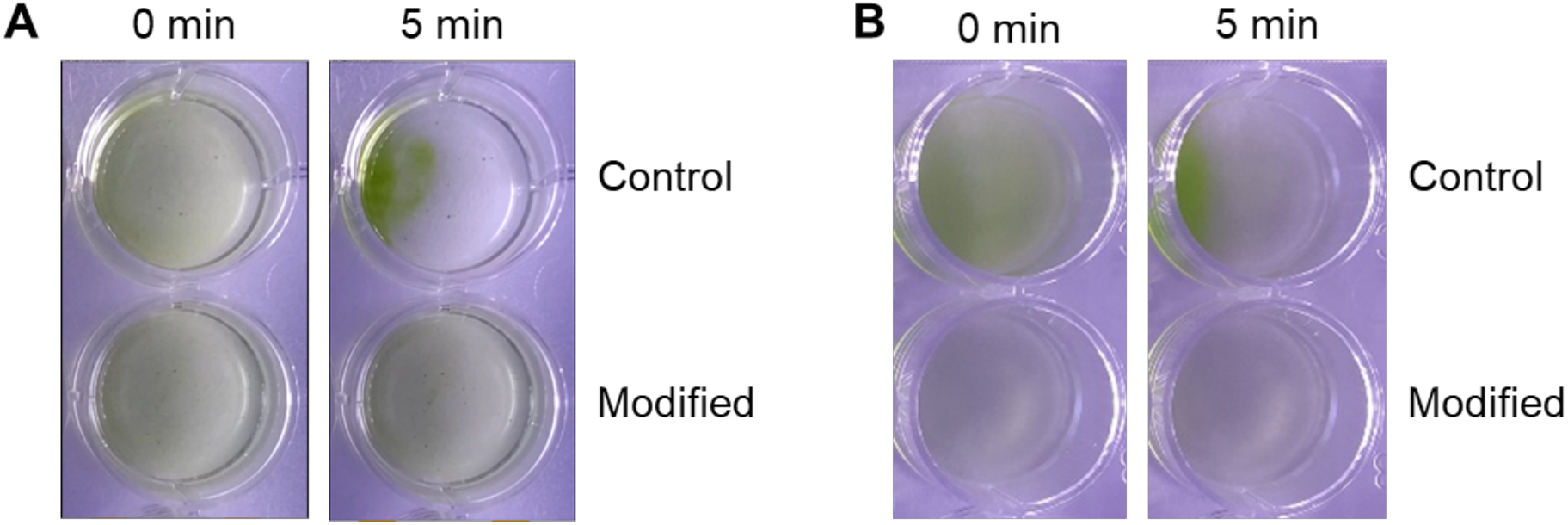
Images of algae before and after visible light illumination. **A**: Control – non-modified microalgae; Modified – microalgae functionalized with DBCO (0.4 mM) and VAN-photoN_3_ (**2**, 0.04 mM). **B**: Control – non-modified microalgae; Modified – microalgae functionalized with DBCO (0.4 mM) and CIP-photoN_3_ (**6**, 0.04 mM).

**Figure S18.**
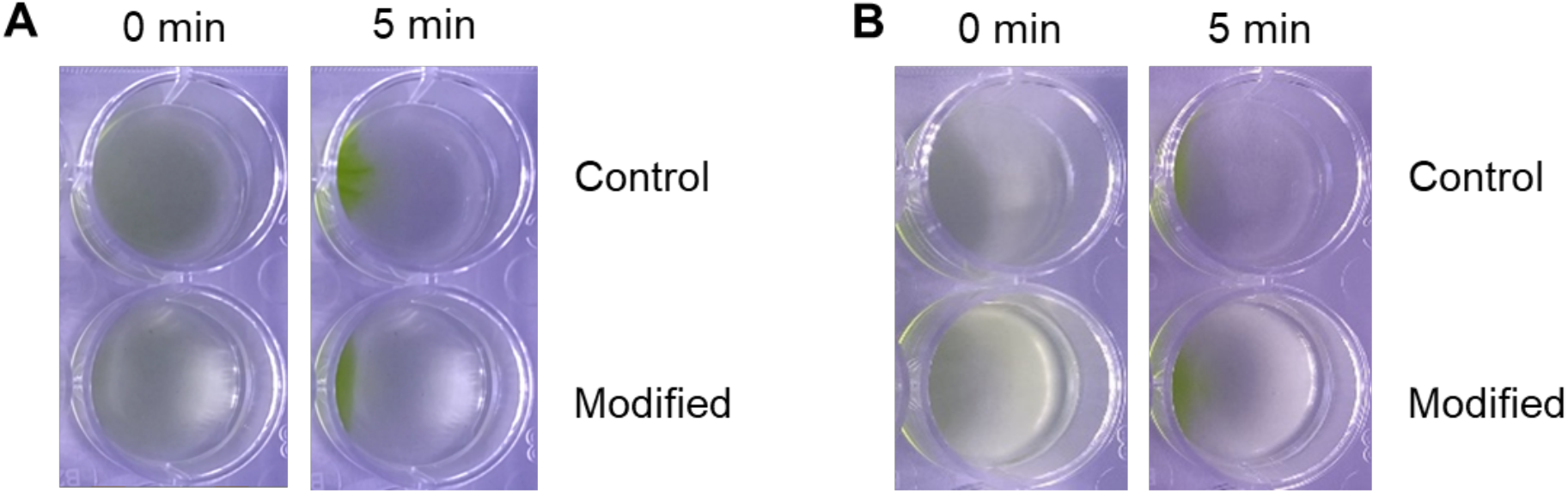
Images of algae before and after visible light illumination. **A**: Control – non-modified microalgae; Modified – microalgae functionalized with DBCO (0.2 mM) and VAN-photoN_3_ (**2**, 0.02 mM). **B**: Control – non-modified microalgae; Modified – microalgae functionalized with DBCO (0.2 mM) and CIP-photoN_3_ (**6**, 0.02 mM).

### 4.3 Drug release studies with and without UV-light after the movement of the biohybrid microswimmers

The modified microalgae collected from left and right side of the wells (section 4.2) were transferred into Eppendorf tubes and kept there for 2 h. Then the cells were centrifuged (3 min, 2000 rcf/min), the supernatant was removed and the cells were washed with dTAP (0.5 mL), centrifuged (3 min, 2000 rcf/min) and the supernatant was removed. The washing step was repeated 6 times. The supernatants after the first centrifugation and after each washing step were collected, combined, and analyzed by UHPLC-MS.

Next, the microalgae were resuspended in dTAP (0.5 mL), loaded in a 24-well plate, and UV irradiated at λ = 365 nm for 5 min. The cells were transferred into Eppendorf tubes, centrifuged (3 min, 2000 rcf/min) and the supernatant was removed. The washing steps were repeated 6 times. The solutions were collected, combined, and analyzed by UHPLC-MS.

**Figure S19.**
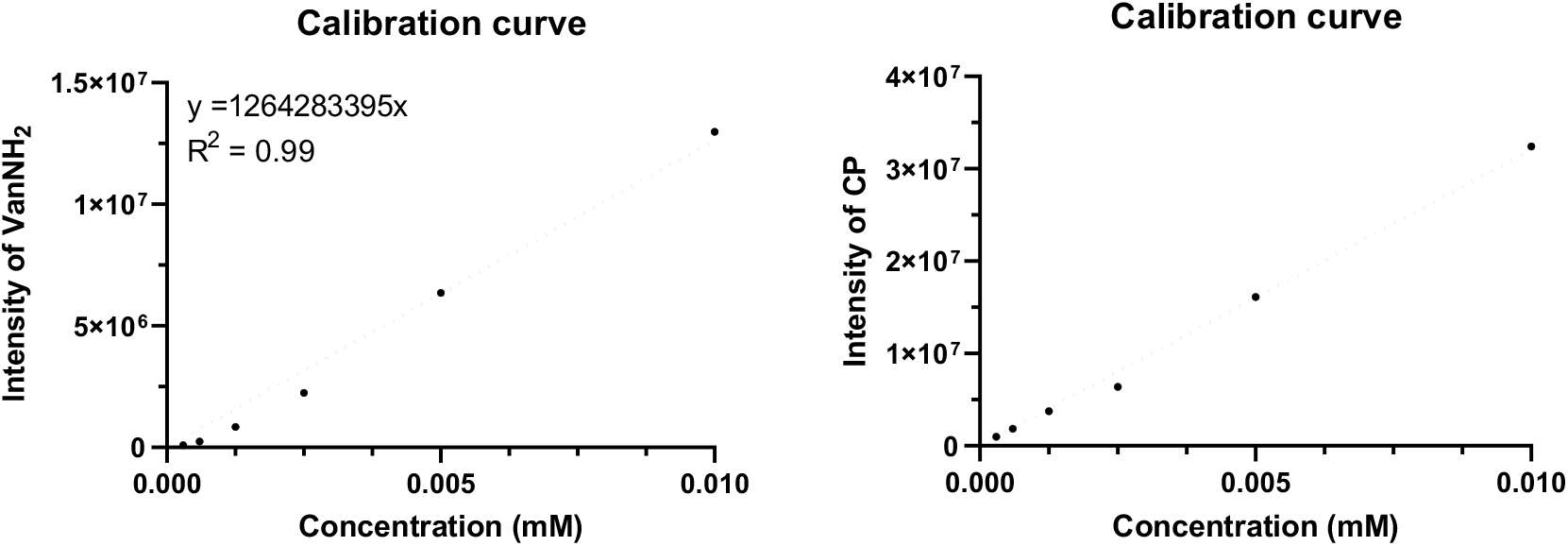
Calibration curves for VAN-NH_2_ (3) on the left and for CIP on the right for the experiment (section 4.3).

## 5. Toxicity experiments

Two-fold serial dilutions of the antibiotics **2** or **6** (ranging from 0.04 mM to 0.0025 mM) were prepared in TAP medium on a 24-well plate. A microalgae culture (0.5 mL) with a cell density 1.5×10^6^ cells/ml was centrifuged (5 min, 4000 rcf/min) and the supernatant was removed. The microalgae were resuspended in fresh TAP medium and the cells were placed into a 24-well plate with final concentration 1.5×10^6^ cells/mL. The microplates were shaken (500 rcf/min) during 4 days at 25 °C and the optical density (OD_750_) was measured by plate reader every day (Synergy H1, BioTek Instruments, VT, USA).

**Figure S20.**
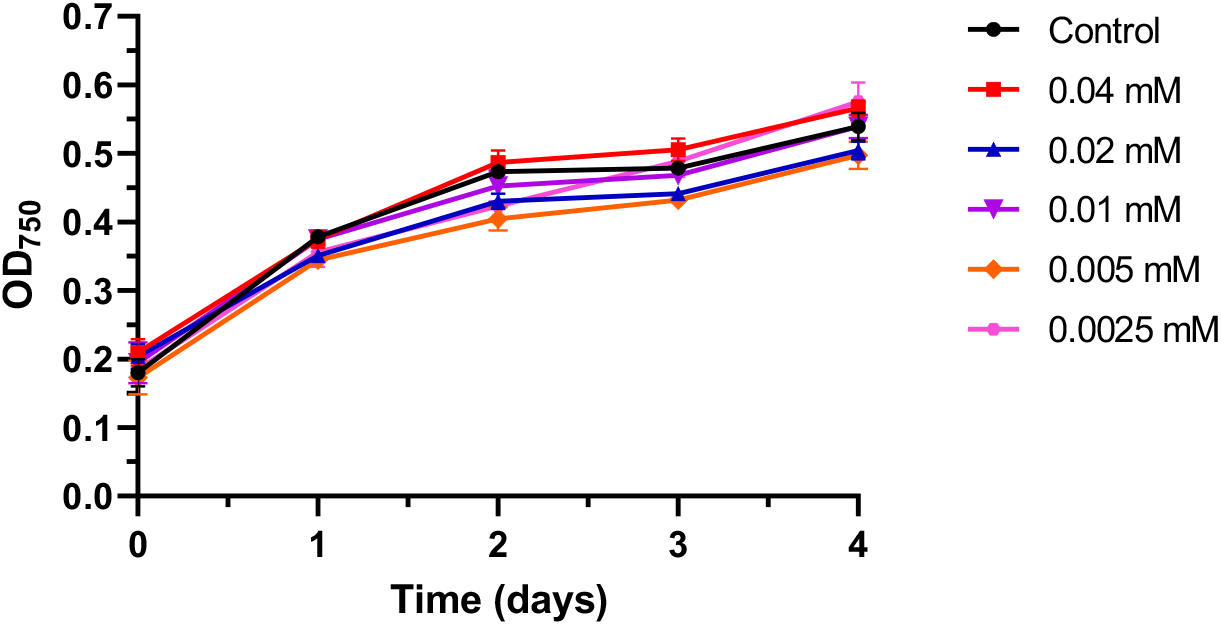
Toxicity experiment of VAN-photoN_3_ (**2**) at different concentrations on microalgae (1.5×10^6^ cells/mL).

**Figure S21.**
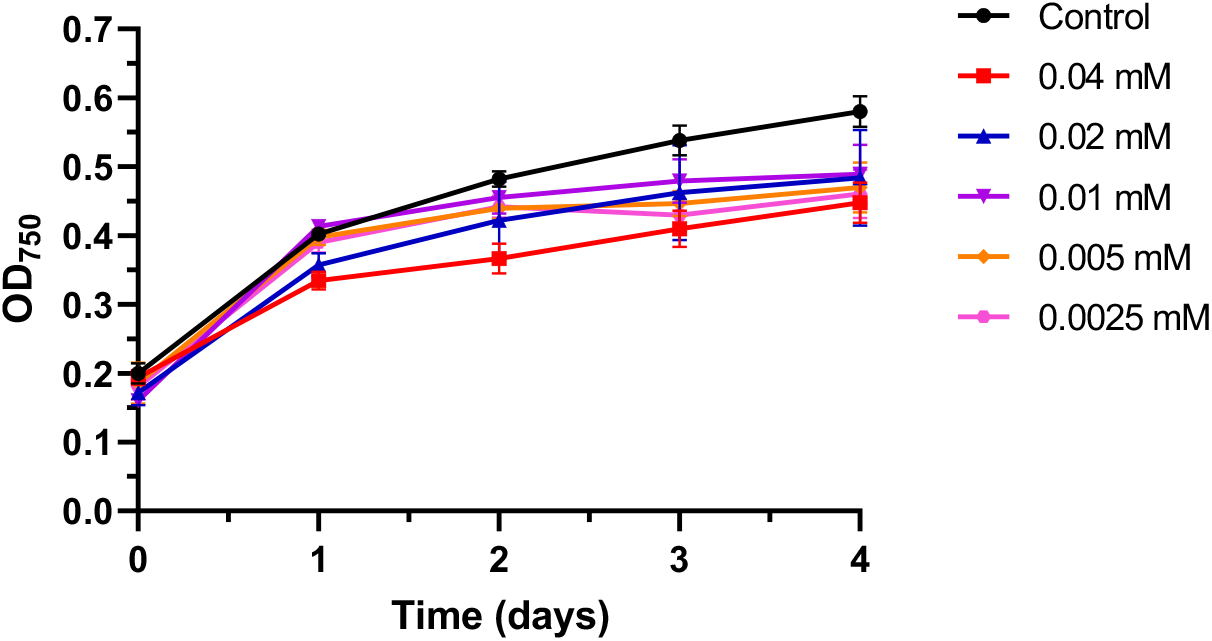
Toxicity experiment of CP-photoN_3_ at different concentrations on algae (1.5×10^6^ cells/mL).

## 6. Light-driven steering of the microalgal swarm

The microalgae (3×10^6^ cells/mL, 2 mL) were placed into cell culture dish (Corning, 35 mm × 10 mm) and the sample was covered to avoid undesired light. The light was shined from the bottom of Petry dish, and the shadow dot was located in a side of the dish for 2 min forcing the cells to cluster in the dark part. Then, the shadow was slowly moved to the other side of the Petry dish over 12 min. The steering of microalgae cluster was recorded and the speed was calculated by the general equation for the speed calculation.

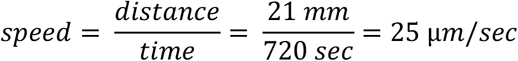

**Figure S22.**
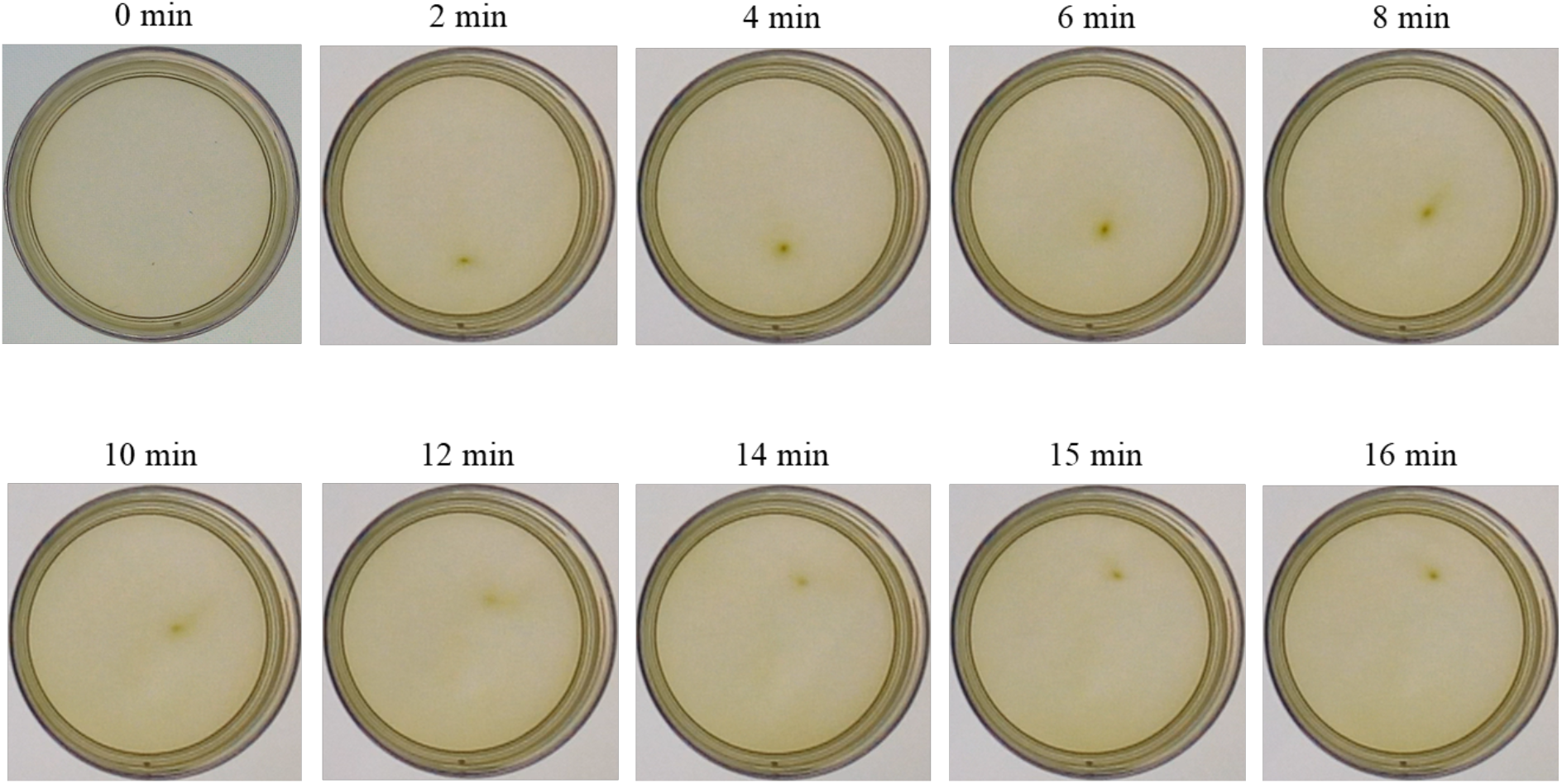
The microalgae swarm movement in the Petry dish (35 mm) during the 16 min of visible light-exposure.

## 7. Antimicrobial activity experiments

### 7.1 MIC value determination

In 96-well microtiter plate, two-fold serial dilutions of the respective antibiotics (ranging from 32 μg/ml to 0.06 μg/ml) were prepared in Mueller-Hinton-II broth (MHB) in a final volume of 50 μl. The bacterial suspensions turbidity was adjusted to a McFarland Standard 0.5 (absorbance at 600 nm 0.08–0.13) to get approximately 1×10^8^ cfu ml^−1^, then the bacterial suspension was diluted by a factor of 1:100 for *S. aureus* ATCC 43300, by a factor 1:200 for *B. subtilis* ATCC 6633 and by a factor 1:150 for *E. coli* ATCC 25922 in MHB. Each well containing the antibiotic solution and the growth control well were inoculated with 50 μl of the bacterial suspension. This results in the final desired inoculum of 5 × 10^5^ cfu ml^−1^ in a volume 100 μl. The plate was then incubated at 37 °C for 18 h, after which minimal inhibitory concentration (MIC) was determined by visual inspection and it is defined as the minimal concentration of a compound that prevents microbial growth.

**Table S2:** MIC value determination of compounds

| Bacteria strain | MIC (µg/ml) |  |  |  |  |
| --- | --- | --- | --- | --- | --- |
|  | VAN | VAN-photoN<sub>3</sub> (**2**) | VAN-NH<sub>2</sub> (**3**) | CIP-photoN<sub>3</sub> (**6**) | CIP |
| *B. subtilis* ATCC 6633 | 0.125-0.25 | 0.25-0.5 | 2-16 | - | - |
| *S. aureus* ATCC 43300 | 1-2 | 1-2 | 1 | 1 | 0.5-1 |
| *E. coli* ATCC 25922 | - | - | - | 0.06 | 0.008 |

### 7.2 Antibacterial activity of the systems

#### 7.2.1 General protocol

The overnight bacterial culture was diluted to an optical density (600 nm) of 0.1 by adding a mixture of biohybrid microalgae in TAP (0.250 ml) and MHB (0.250 ml). The mixture was UV-irradiated at λ=365 nm for 5 min (unless stated otherwise) and grown in a 24-well plate at 37 °C. The cell density (600 nm) was measured every 15 min for 18 h (with shaking between measurements) in a microplate reader. All experiments were performed in triplicates.

#### 7.2.2 Low concentration protocol

The microalgae were exposed to the same incubation and washing steps as was described in the experimental section in the manuscript, *Preparation of the biohybrids* section, DBCO (0.05 mM) was added during the 1^st^ step and CIP-photoN_3_ (**6**, 0.0025 mM or 0.005 mM) was added during the 2^nd^ step. The antibacterial activity of the system was recorded by replacing the TAP medium (0.250 mL) from the general protocol **7.2.1**. with the microalgae resuspended in fresh TAP Medium (0.250 mL). The bacterial growth was recorded with or without the irradiation step.

**Figure S23.**
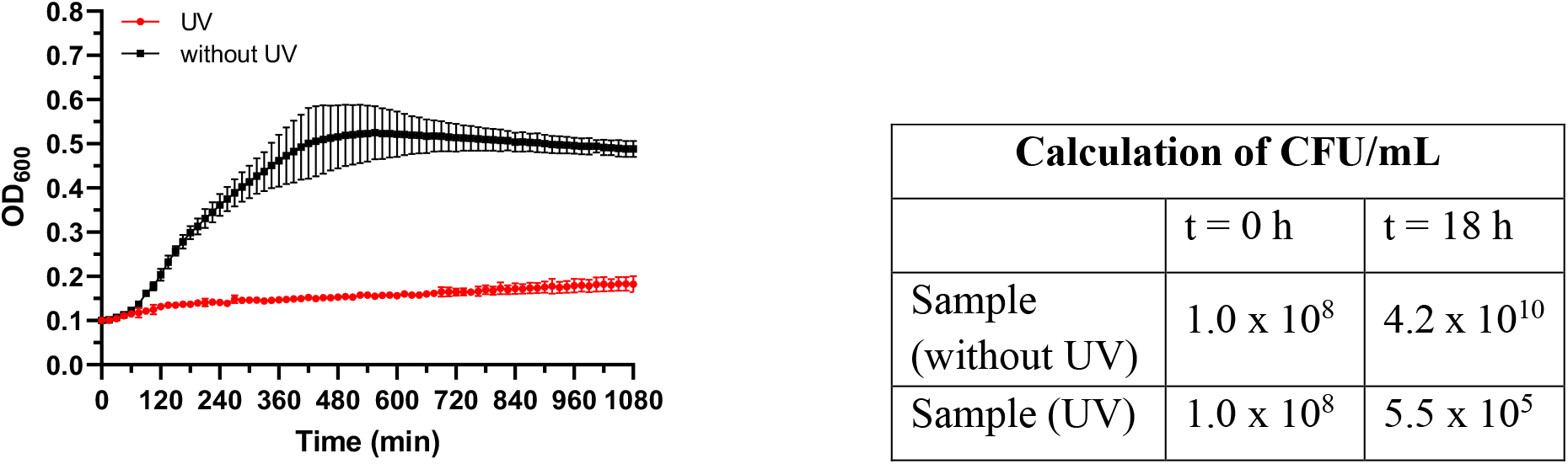
Growth of *E. coli* incubated with microalgae functionalized with DBCO (0.05 mM) and CIP-photoN_3_ (**6**, 0.0025 mM). Left: Non-irradiated samples – black line; the samples irradiated at time 0 for 5 min (λ = 365 nm) – red line. Right: CFU calculation of bacteria from the samples at time 0 min and after 18 h of bacterial growth. Data points represent mean value ± SD (n=3).

**Figure S24.**
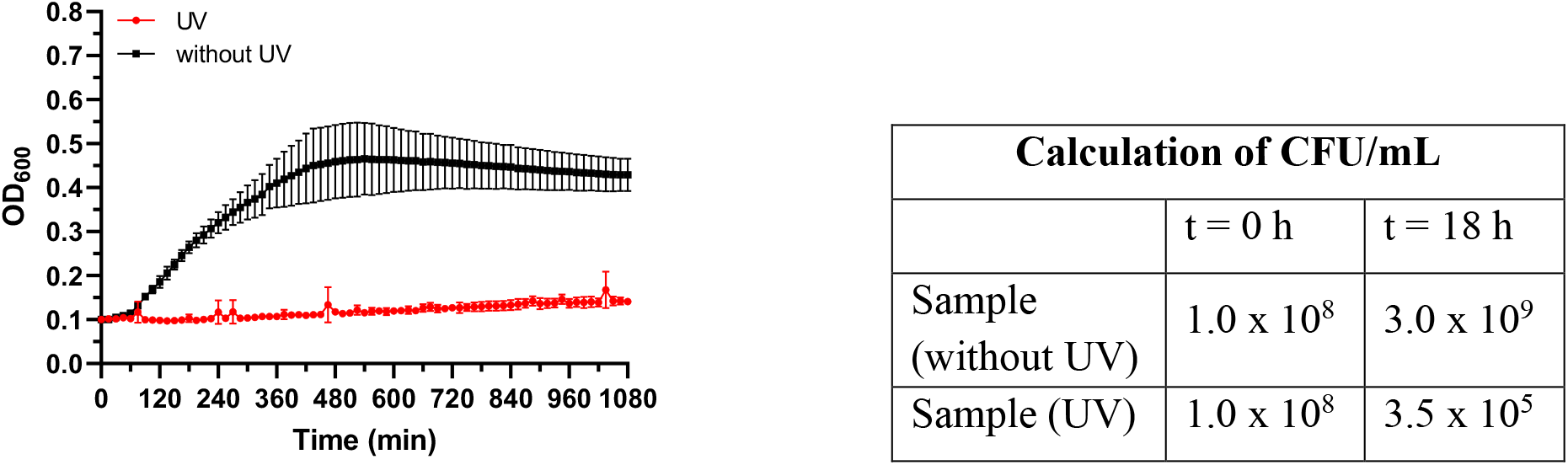
Growth of *E. coli* incubated with microalgae functionalized with DBCO (0.1 mM) and CIP-photoN_3_ (**6**, 0.005 mM). Left: Non-irradiated samples – black line; the samples irradiated at time 0 for 5 min (λ = 365 nm) – red line. Right: CFU calculation of bacteria from the samples at time 0 min and after 18 h of bacterial growth. Data points represent mean value ± SD (n=3).

#### 7.2.3 High concentration protocol

The microalgae were exposed to the same incubation and washing steps as was described in the experimental section in the manuscript, *Preparation of the biohybrids* section, DBCO (0.02 mM) was added during the 1^st^ step and CIP-photoN_3_ or VAN-photoN_3_ (**6** or **2**, 0.2 mM) was added during the 2^nd^ step. The antibacterial activity of the system was recorded by replacing the TAP medium (0.250 mL) from the general protocol **7.2.1**. with the microalgae resuspended in fresh TAP Medium (0.250 mL). The bacterial growth was recorded with or without the irradiation step. For the testing against *S. aureus* two samples of modified microalgae was combined and resuspended in fresh TAP medium (0.250 mL).

**Figure S25.**
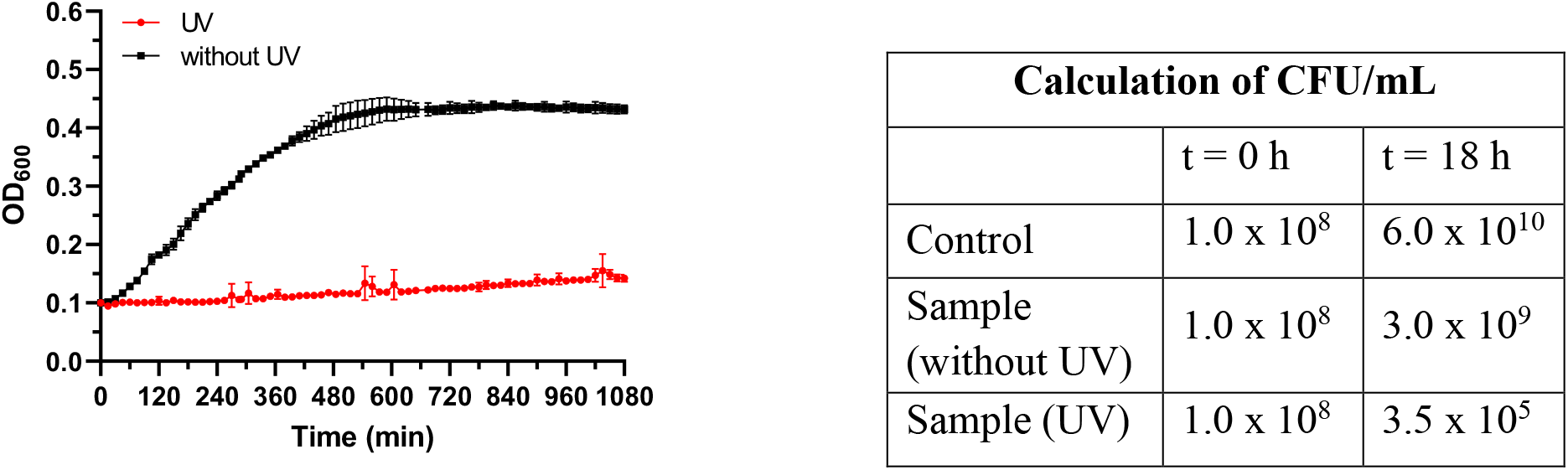
Growth of *E. coli* incubated with algae functionalized with DBCO (0.2 mM) and CIP-photoN_3_ (**6**, 0.02 mM). Left: Non-irradiated samples – black line; the samples irradiated at time 0 for 5 min (λ = 365 nm) – red line. Right: CFU calculation of bacteria from the samples at time 0 min and after 18 h of bacterial growth. Data points represent mean value ± SD (n=3).

**Figure S26.**
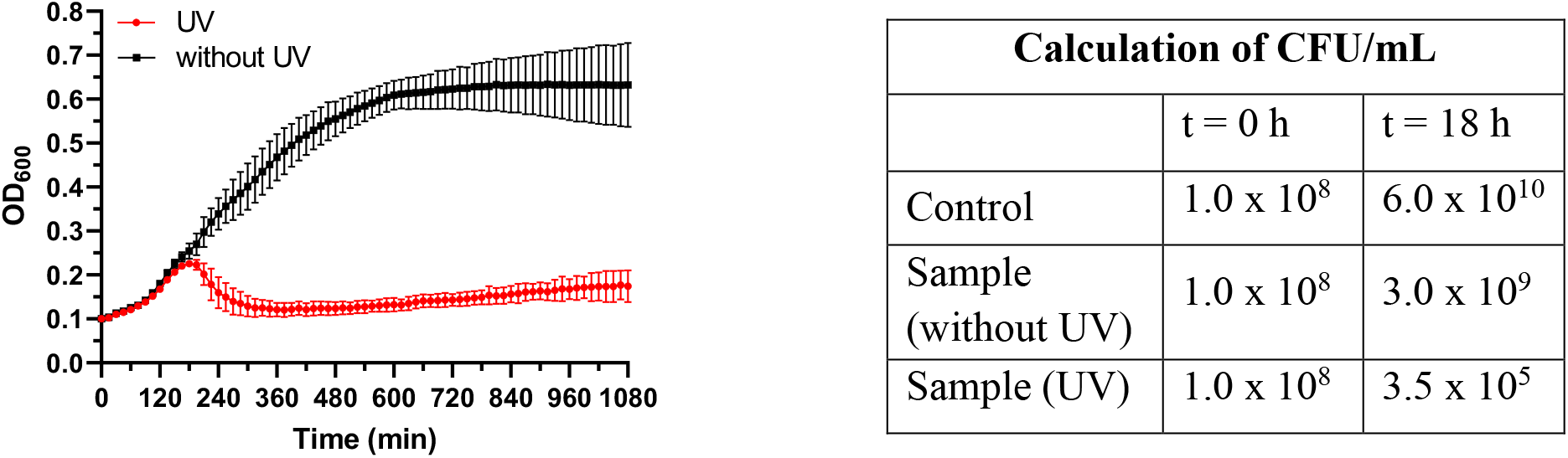
Growth of *S. aureus* incubated with microalgae functionalized with DBCO (0.2 mM) and CIP-photoN_3_ (**6**, 0.02 mM). Two samples of modified microalgae were combined together for testing. Left: Non-irradiated samples – black line; the samples irradiated at time 0 for 5 min (λ = 365 nm) – red line. Right: CFU calculation of bacteria from the samples at time 0 min and after 18 h of bacterial growth. Data points represent mean value ± SD (n=3).

**Figure S27.**
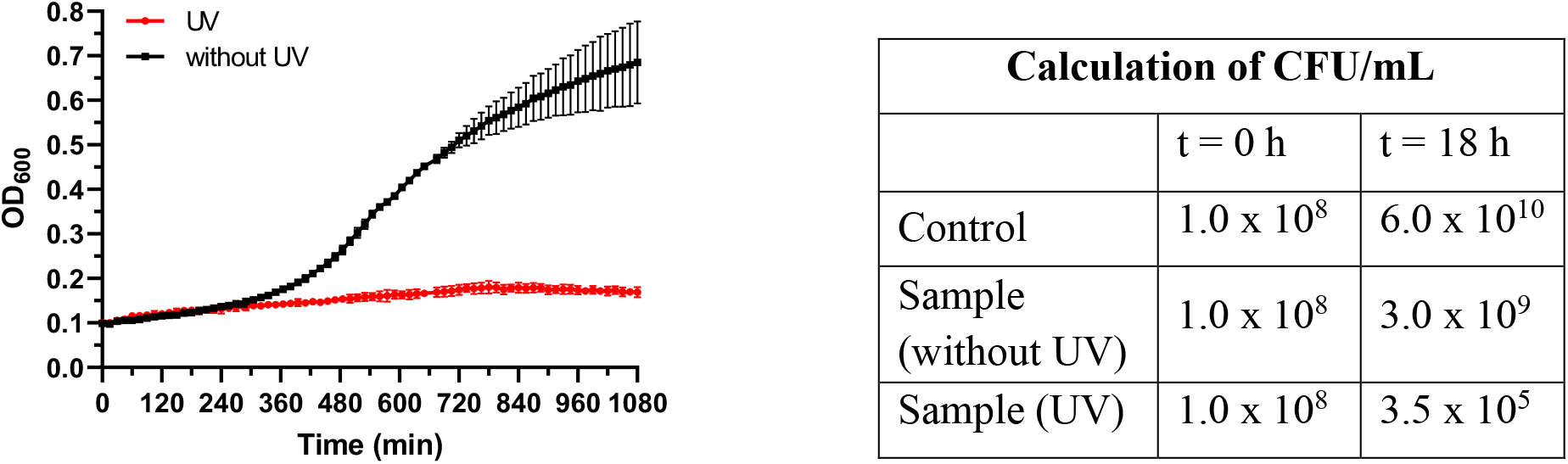
Growth of *S. aureus* incubated with microalgae functionalized with DBCO (0.2 mM) and VAN-photoN_3_ (**2**, 0.02 mM). Two samples of modified microalgae were combined together for testing. Left: Non-irradiated samples – black line; the samples irradiated at time 0 for 5 min (λ = 365 nm) – red line. Right: CFU calculation of bacteria from the samples at time 0 min and after 18 h of bacterial growth. Data points represent mean value ± SD (n=3).

### 7.3 Control experiments

#### 7.3.1 Antibacterial activity of C. reinhardtii functionalized with DBCO

The algae were exposed to the same incubation and washing steps as was described in the experimental section in the manuscript, *Preparation of the biohybrids* section, DBCO (0.2 mM) was added during the 1^st^ step and either VAN-photoN_3_ (**2**) or CIP-photoN_3_ (**6**) were not added during the 2^d^ step. The antibacterial activity of the system was recorded by replacing the Kuhl medium (0.250 mL) from the general protocol **7.2.1**. with the microalgae resuspended in fresh Kuhl Medium (0.250 mL). For the testing against *S. aureus* two samples of modified microalgae were combined and resuspended in fresh TAP medium (0.250 mL).

**Figure S28.**
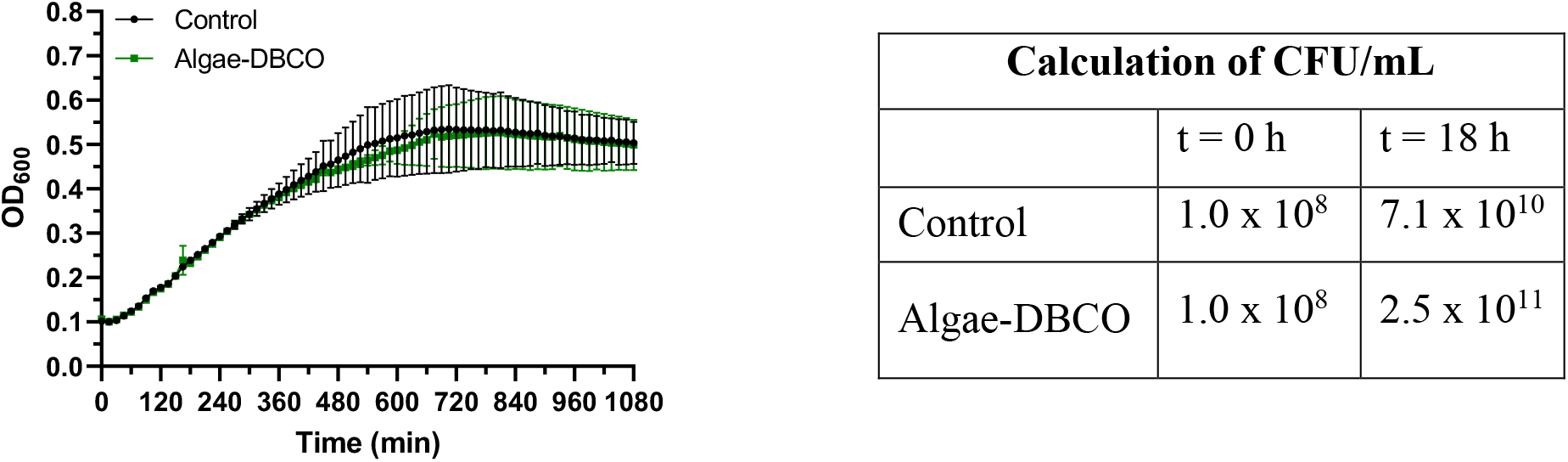
Growth of *E. coli* incubated with microalgae functionalized with DBCO (0.02 mM) and after the 5 min irradiation step (λ = 365 nm). Left: bacterial growth in presence of non-modified microalgae – black line; bacterial growth in presence of microalgae modified with DBCO (0.02 mM) and irradiated at time 0 for 5 min (λ = 365 nm) – green line. Right: CFU calculation of bacteria from the samples at time 0 min and after 18 h of bacterial growth. Data points represent mean value ± SD (n=3).

**Figure S29.**
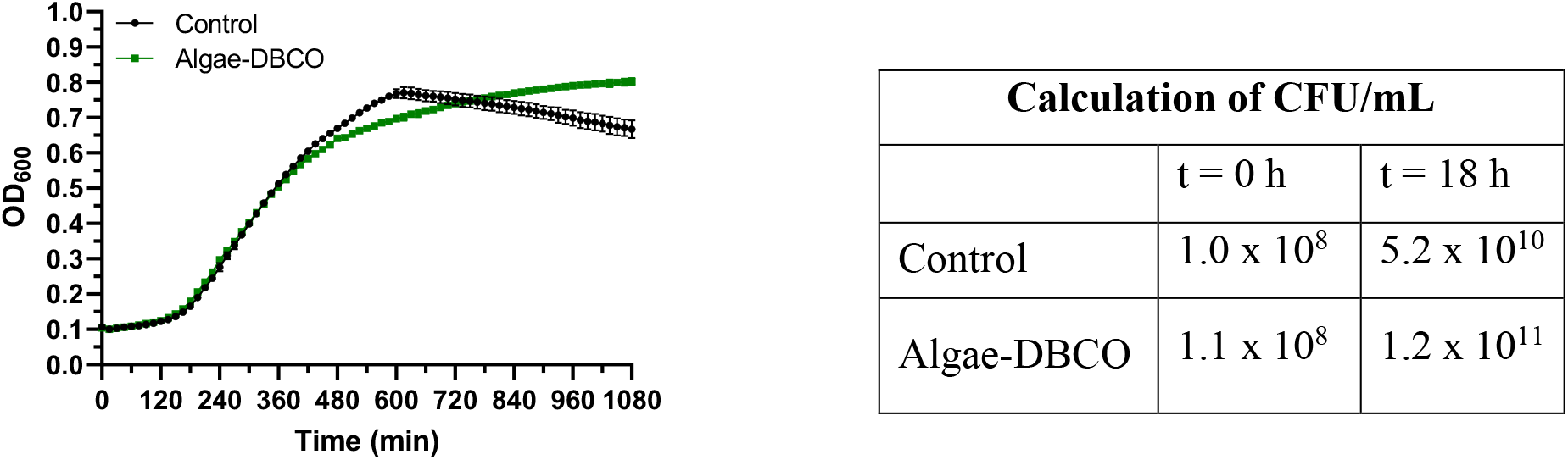
Growth of *S. aureus* incubated with microalgae functionalized with DBCO (0.02 mM) and UV-irradiated (λ = 365 nm) for 5 min. Left: bacterial growth in presence of non-modified microalgae – black line; bacterial growth in presence of microalgae modified with DBCO (0.02 mM) and irradiated at time 0 for 5 min (λ = 365 nm) – green line. Right: CFU calculation of bacteria from the samples at time 0 min and after 18 h of bacterial growth. Data points represent mean value ± SD (n=3).

#### 7.3.2 Antibacterial activity of C. reinhardtii only treated with VAN-photoN_3_

The algae were exposed to the same incubation and washing steps as was described in the experimental section in the manuscript, *Preparation of the biohybrids* section, DBCO was not added during the 1^st^ step and VAN-photoN_3_ (**2**, 0.02 mM) were added during the 2^nd^ step. The antibacterial activity of the system was recorded by replacing the TAP medium (0.250 mL) from the general protocol **7.2.1**. with the microalgae resuspended in fresh TAP Medium (0.250 mL). The bacterial growth was recorded with or without the irradiation step. Two samples of modified microalgae were combined and resuspended in fresh TAP medium (0.250 mL).

**Figure S30.**
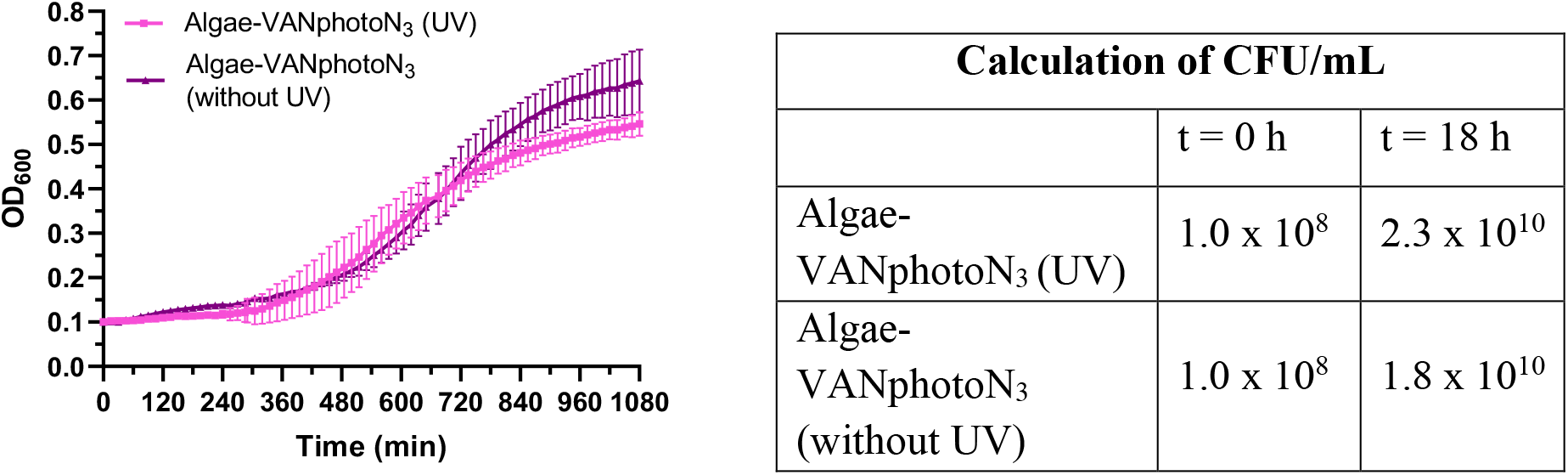
Growth of *S. aureus* incubated with microalgae treated with VAN-photoN_3_ (**2**, 0.02 mM). Left: bacterial growth in presence of modified microalgae UV-irradiated (λ = 365 nm) for 5 min and not exposed to UV-light. Right: CFU calculation of bacteria from the samples at time 0 min and after 18 h of bacterial growth. Data points represent mean value ± SD (n=3)

#### 7.3.3 Antibacterial activity of C. reinhardtii only treated with CIP-photoN_3_

The microalgae were exposed to the same incubation and washing steps as was described in the experimental section in the manuscript, *Preparation of the biohybrids* section, DBCO was not added during the 1^st^ step and CIP-photoN_3_ (**6**, 0.02 mM) were added during the 2^nd^ step. The antibacterial activity of the system was recorded by replacing the TAP medium (0.250 mL) from the general protocol **7.2.1**. with the microalgae resuspended in fresh TAP Medium (0.250 mL). The bacterial growth was recorded with or without the irradiation step. For the testing against *S. aureus* two samples of modified microalgae were combined and resuspended in fresh TAP medium (0.250 mL).

**Figure S31.**
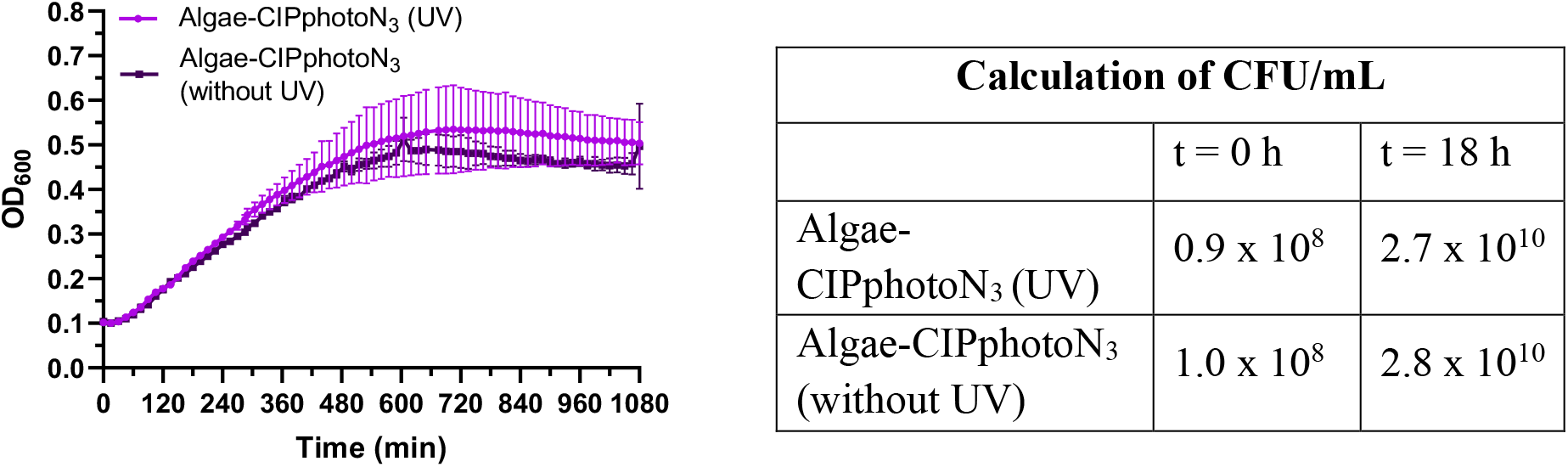
Growth of *E. coli* incubated with microalgae treated with CIP-photoN_3_ (**6**, 0.02 mM). Left: bacterial growth in presence of modified microalgae UV-irradiated (λ = 365 nm) for 5 min and not exposed to UV-light. Right: CFU calculation of bacteria from the samples at time 0 min and after 18 h of bacterial growth. Data points represent mean value ± SD (n=3)

**Figure S32.**
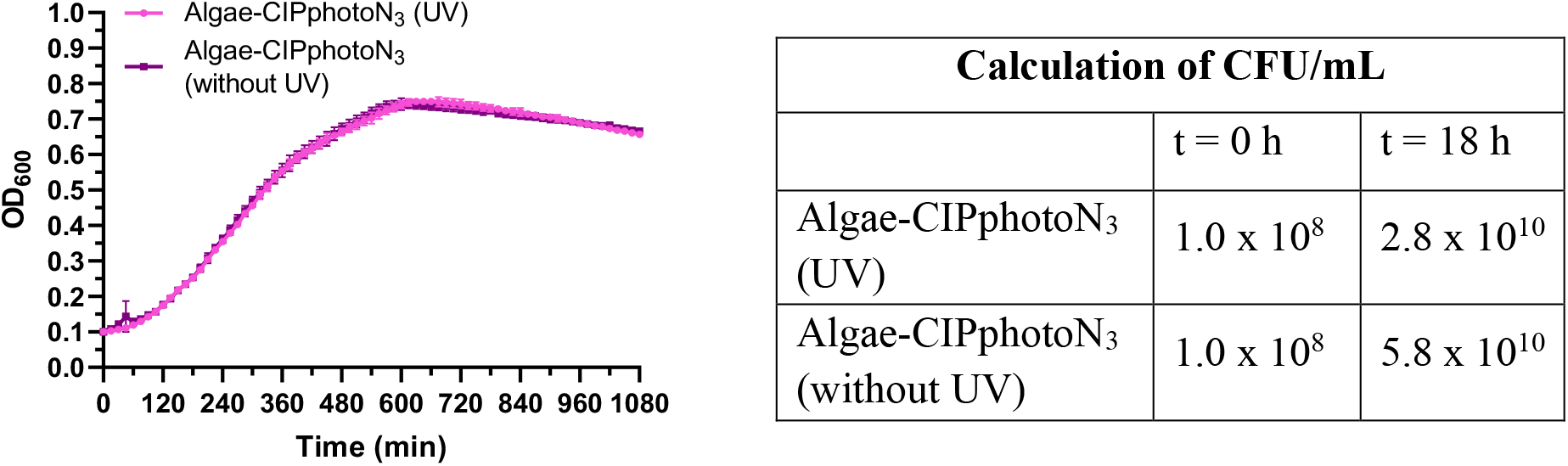
Growth of *S. aureus* incubated with microalgae treated with CIP-photoN_3_ (**6**, 0.02 mM). Left: bacterial growth in presence of modified microalgae UV-irradiated (λ = 365 nm) for 5 min and not exposed to UV-light. Right: CFU calculation of bacteria from the samples at time 0 min and after 18 h of bacterial growth. Data points represent mean value ± SD (n=3)

#### 7.3.4 Antibacterial activity of the functionalized C. reinhardtii after the irradiation step

The microalgae were exposed to the same incubation and washing steps as was described in the experimental section in the manuscript, *Preparation of the biohybrids* section. The functionalised *C. reinhardtii* cells were resuspended in dTAP (0.5 mL), transferred into the 24-well plate and UV-irradiated at λ = 365 nm for 5 min. Released VAN-NH_2_ or CIP was washed out with dTAP (1 mL), the sample was centrifuged (3 min, 2000 rcf/min), the supernatant was removed and the washing step was repeated 6 times. The antibacterial activity of the system was recorded by replacing the TAP medium (0.250 mL) from the general protocol **7.2.1**. with the algae resuspended in fresh TAP medium (0.250 mL). For the testing against *S. aureus* two samples of modified microalgae were combined and resuspended in fresh TAP medium (0.250 mL).

**Figure S33.**
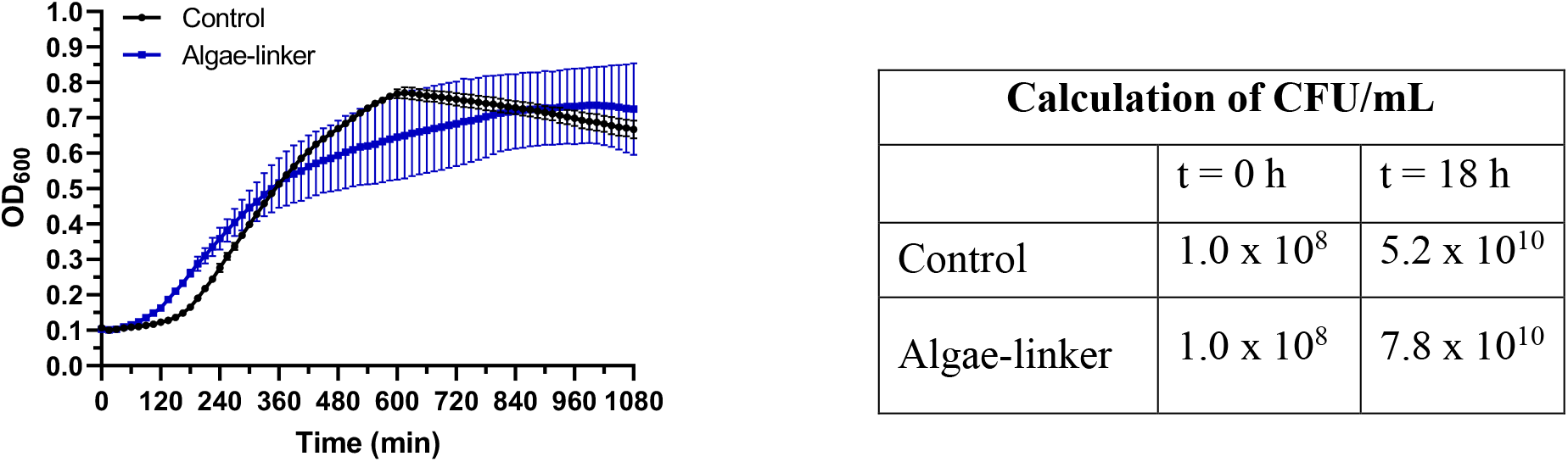
Growth of *S. aureus* incubated with the functionalized microalgae after the UV-irradiation step (λ = 365 nm). Left: bacterial growth in presence of non-modified microalgae – black line; bacterial growth in presence of modified microalgae with DBCO (0.02 mM) and CIP-photoN_3_ (**6**, 0.02 mM) and UV-irradiated (λ = 365 nm) – blue line. Right: CFU calculation of bacteria from the samples at time 0 min and after 18 h of bacterial growth. Data points represent mean value ± SD (n=3).

**Figure S34.**
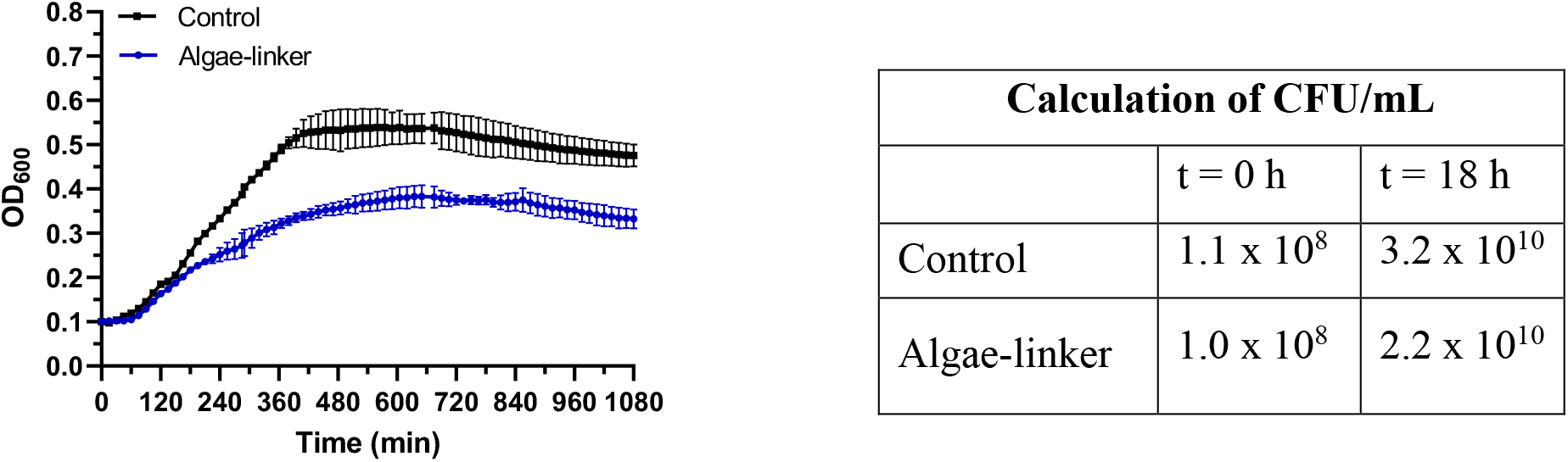
Growth of *E. coli* incubated with the functionalized microalgae after the UV-irradiation step (λ = 365 nm). Left: bacterial growth in presence of non-modified microalgae – black line; bacterial growth in presence of modified microalgae with DBCO (0.02 mM) and CIP-photoN_3_ (**6**, 0.02 mM) and UV-irradiated (λ = 365 nm) – blue line. Right: CFU calculation of bacteria from the samples at time 0 min and after 18 h of bacterial growth. Data points represent mean value ± SD (n=3).

**Figure S35.**
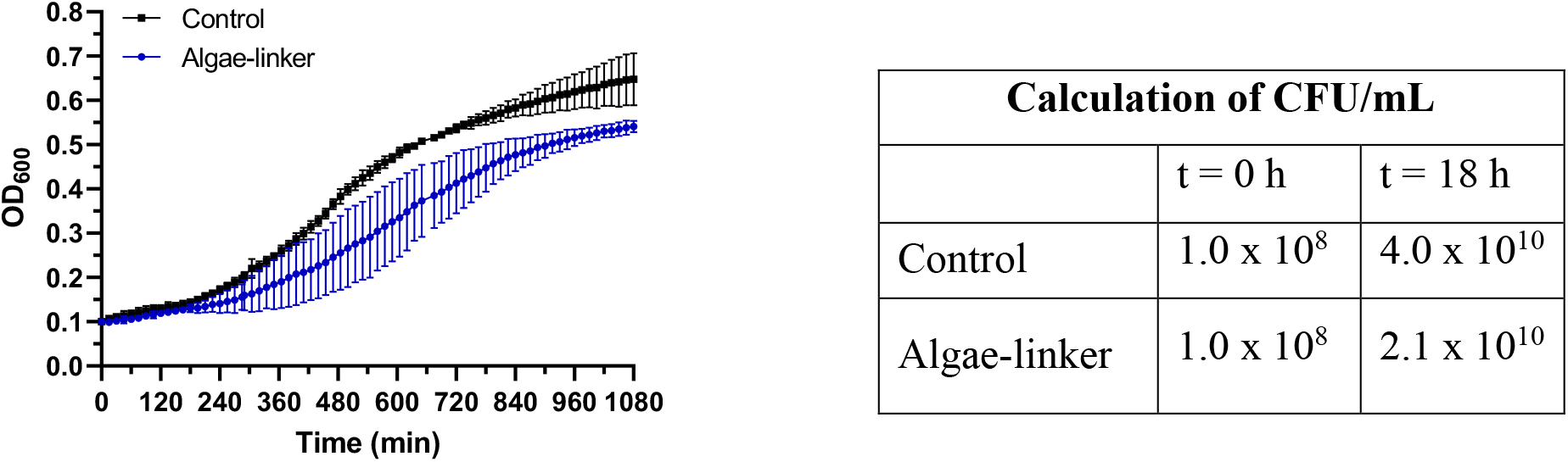
Growth of *S. aureus* incubated with the functionalized microalgae after the UV-irradiation step (λ = 365 nm). Left: bacterial growth in presence of non-modified microalgae – black line; bacterial growth in presence of modified microalgae with DBCO (0.02 mM) and VAN-photoN_3_ (**2**, 0.02 mM) and UV-irradiated (λ = 365 nm) – blue line. Right: CFU calculation of bacteria from the samples at time 0 min and after 18 h of bacterial growth. Data points represent mean value ± SD (n=3).

#### 7.3.5 Influence of UV-irradiation on bacterial growth

To determine the influence of the irradiation time on the viability of *E. coli* and *S*. *aureus*, bacterial solutions (OD_600_ = 0.1) were irradiated for 0 or 5 min at λ = 365 nm and the culture growths were recorded.

**Figure S36.**
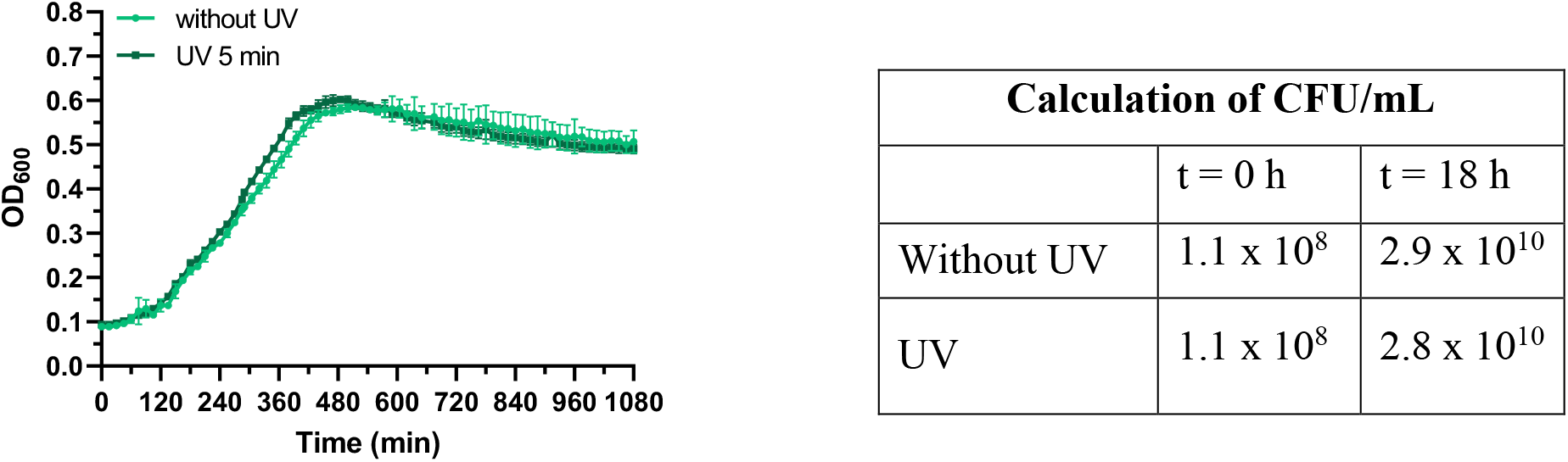
Growth of *E. coli* after the UV-irradiation step (λ = 365 nm) during 0 or 5 min. Left: bacterial growth without UV-irradiation – light green line; bacterial growth after UV-irradiation (λ = 365 nm) – dark green line. Right: CFU calculation of bacteria from the samples at time 0 min and after 18 h of bacterial growth. Data points represent mean value ± SD (n=3).

**Figure S37.**
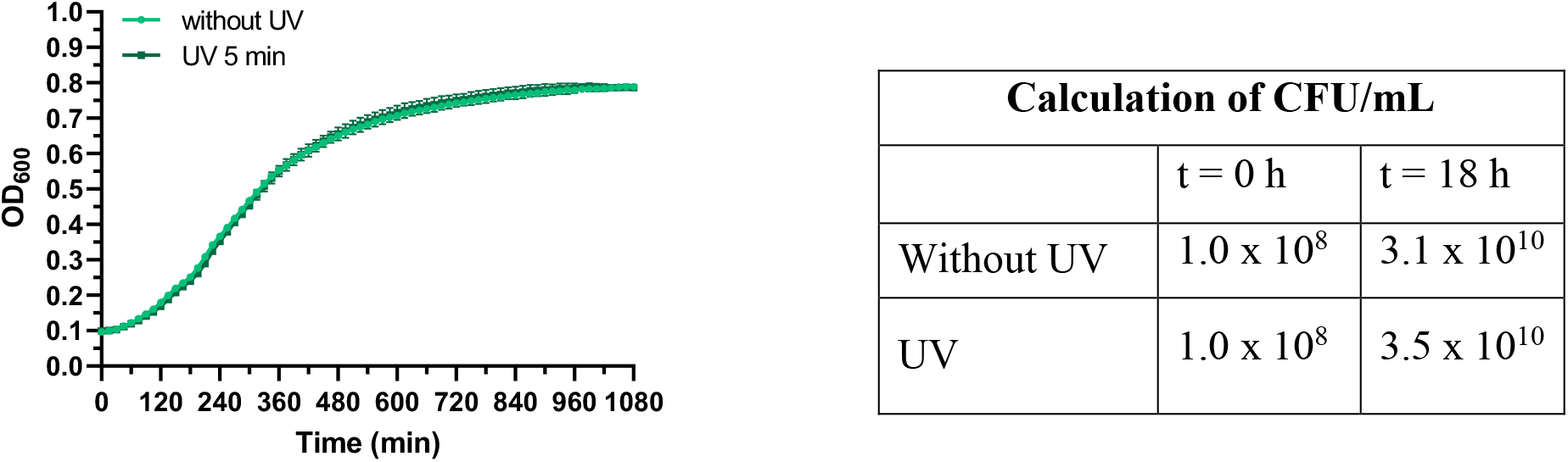
Growth of *S. aureus* after the UV-irradiation step (λ = 365 nm) during 0 or 5 min. Left: bacterial growth without UV-irradiation – light green line; bacterial after UV-irradiation (λ = 365 nm) – dark green line. Right: CFU calculation of bacteria from the samples at time 0 min and after 18 h of bacterial growth. Data points represent mean value ± SD (n=3).

#### 7.3.6 Influence of TAP medium on bacterial growth

To determine the influence of the presence of TAP medium on bacterial growth of *E. coli* and *S*. *aureus*, bacterial solutions (OD_600_ = 0.1) were grown in presence of mixture TAP:MHB 1:1 and MHB and the culture growths was recorded.

**Figure S38.**
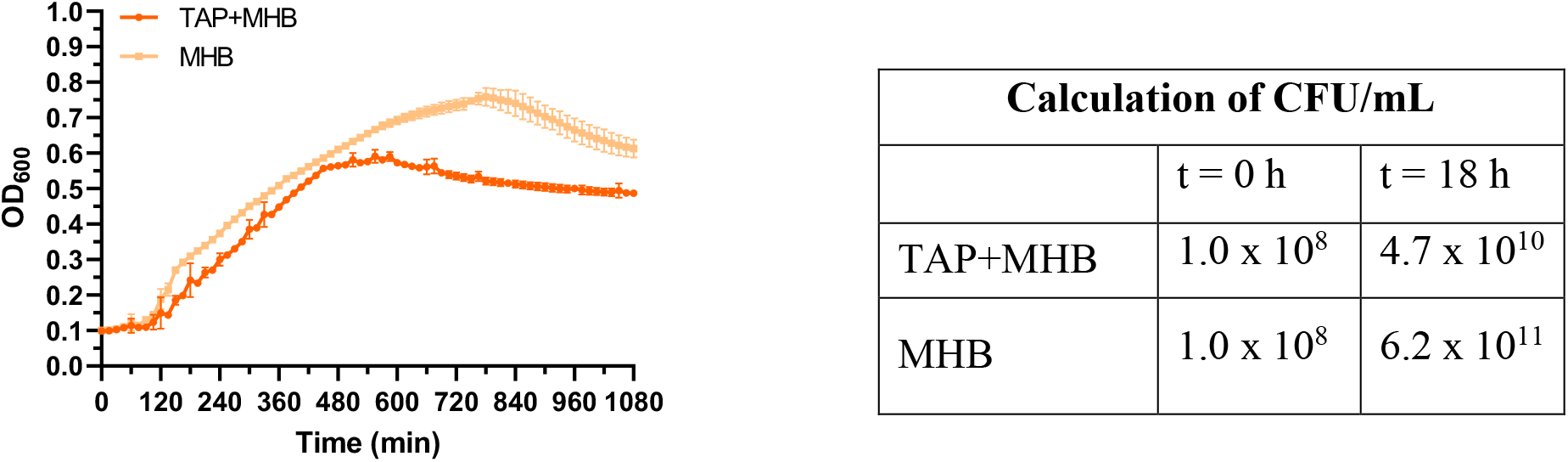
Growth of *E. coli* grown in different mediums. Left: bacterial growth in medium mixture TAP+MHB – dark orange line; bacterial growth in MHB medium – light orange line. Right: CFU calculation of bacteria from the samples at time 0 min and after 18 h of bacterial growth. Data points represent mean value ± SD (n=3).

**Figure S39.**
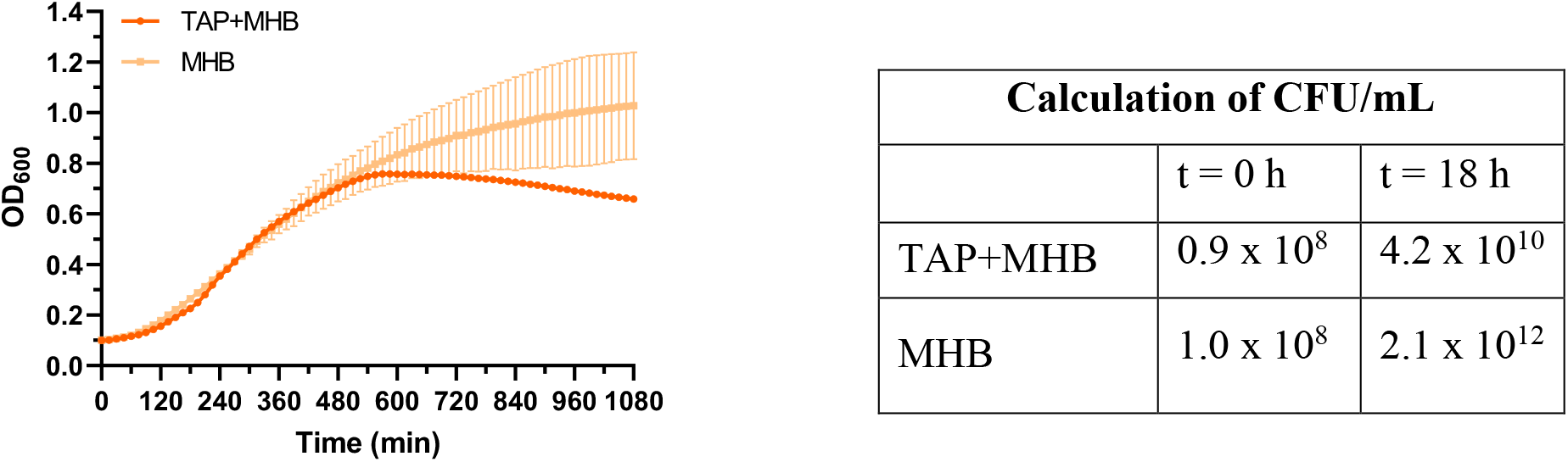
Growth of *S. aureus* grown in different mediums. Left: bacterial growth in medium mixture TAP+MHB – dark orange line; bacterial growth in MHB medium – light orange line. Right: CFU calculation of bacteria from the samples at time 0 min and after 18 h of bacterial growth. Data points represent mean value ± SD (n=3).

#### 7.3.7 Antibacterial activity of the system at exponential phase of bacterial growth

The microalgae were exposed to the same incubation and washing steps as was described in the experimental section in the manuscript, <u>*Preparation of the biohybrids*</u> section), DBCO was added during the 1^st^ step and CIP-photoN_3_ (**6**) was added during the 2^nd^ step according to low concentration protocol 7**.2.2**. The antibacterial activity of the system was recorded by replacing the TAP medium (0.250 mL) from the general protocol 7**.2.1**. with the microalgae resuspended in fresh TAP medium (0.250 mL). The bacterial growth was recorded with the irradiation step after 5 h of bacterial growth.

**Figure S40.**
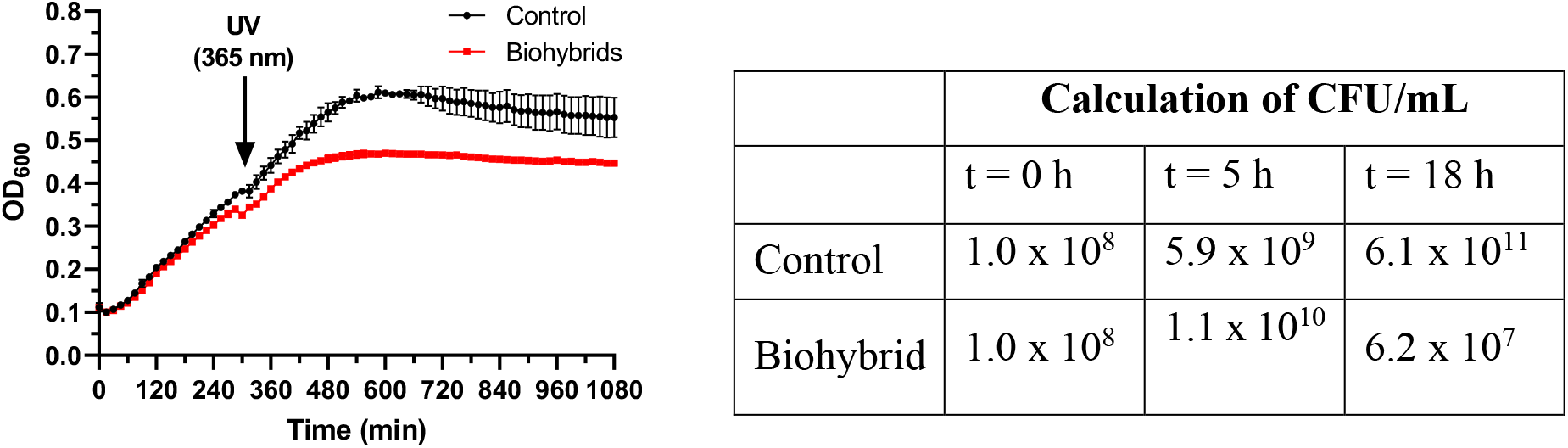
Growth of *E. coli* incubated with biohybrids functionalized with low concentration protocol (DBCO (0.05 mM)) and CIP-photoN_3_ (**6**, 0.005 mM)). The irradiation step (5 min, 365 nm) was applied after 5 h of bacterial growth. Data points represent mean value ± SD (n=3)

## 8. Drug delivery experiment

### 8.1 Experiment with modified algae without visible light exposure

The modified microalgae (0.050 μL) were carefully placed on the right side of the channel. The plate was positioned in the dark and left there for 15 min. The microalgae remained in the channel was carefully removed and the plates were UV-irradiated for 5 min at λ = 365 nm. The plates were next incubated for 18 h at 37 °C. The bacterial growth was determined by visual inspection.

*E. coli* ATCC 25922

**Figure S41.**
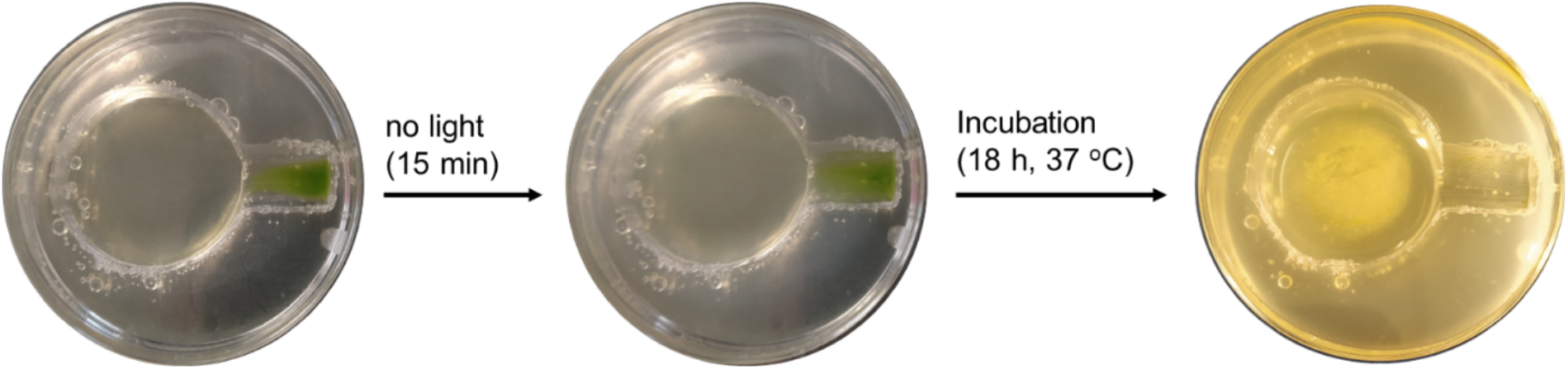
Control experiment with biohybrids. The microalgae were modified according to the general procedure (section 7.2.1; DBCO = 0.2 mM, CIP-photoN_3_ = 0.02 mM) and tested against *E. coli* ATCC 25922.

**Figure S42.**
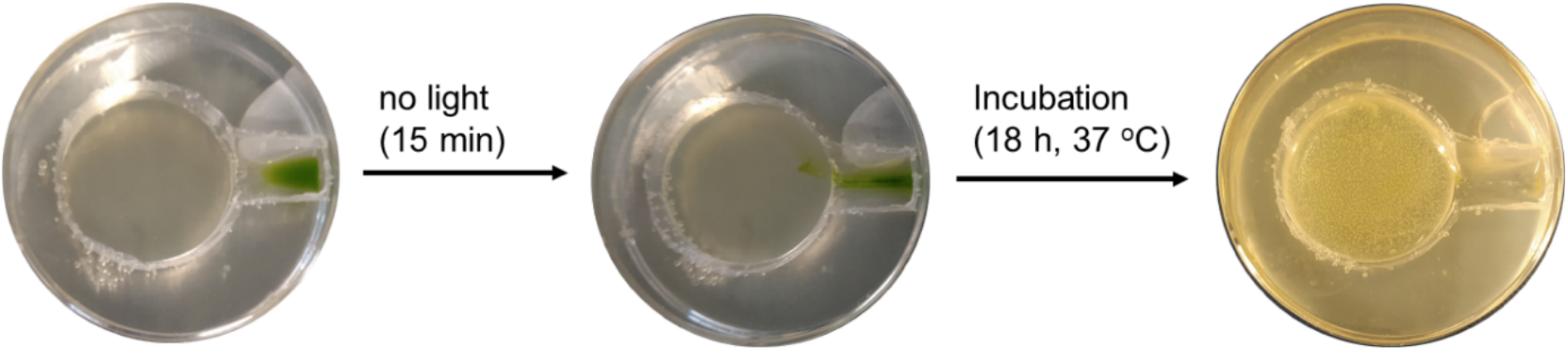
Control experiment with biohybrids. The microalgae were modified according to the general procedure (section 7.2.1; DBCO = 0.2 mM, VAN-photoN_3_ = 0.02 mM) and tested against *B. subtilis* ATCC 6633.

